# CD40 provides immune privilege to the bone marrow hematopoietic niche

**DOI:** 10.1101/2020.08.10.243691

**Authors:** Barbara Bassani, Alessandro Gulino, Paola Portararo, Laura Botti, Barbara Cappetti, Claudia Chiodoni, Niccolò Bolli, Marilena Ciciarello, Korinna Joehrens, Ioannis Anagnostopoulos, Il-Kang Na, Antonio Curti, Claudio Tripodo, Mario P. Colombo, Sabina Sangaletti

**Affiliations:** Department of Research, Fondazione IRCCS Istituto Nazionale Tumori, Milan, Italy; University of Palermo, Palermo, Italy; Department of Oncology and Hemato-Oncology, University of Milan, Milan, Italy; Department of Oncology and Hematology, Fondazione IRCCS Istituto Nazionale dei Tumori, Milan, Italy; Department of Experimental, Diagnostic and Specialty Medicine – DIMES, Institute of Hematology “Seràgnoli”, Bologna, Italy; Institute of Pathology, Charité-Universitätsmedizin Berlin, 10117, Berlin, Germany; Department of Hematology, Oncology and Tumor Immunology, Charité - Universitätsmedizin Berlin, corporate member of Freie Universität Berlin, Humboldt-Universität zu Berlin, and Berlin Institute of Health, Berlin, Germany; Experimental and Clinical Research Center, Berlin, Germany; Berlin-Brandenburg Center for Regenerative Therapies, Berlin, Germany; Berlin Institute of Health, Berlin, Germany

## Abstract

Allogeneic bone marrow transplantation remains the only therapeutic option for a wide range of hematological malignancies despite the risk of possible adverse, immune-related events, such as infection and acute graft-versus-host disease (aGVHD). aGVHD is characterized by T-cell activation, defective B-cell development and osteoblastic niche destruction in bone marrow (BM) among other issues. Transplant conditioning regimens cause excessive inflammatory cytokines production and impaired regulatory T-cell control of aberrant T-cell activation. Here, we show that mesenchymal cells (MSCs) upregulated CD40 upon irradiation at the expense of mesenchymal markers, and that CD40 endows MSC of regulatory function on Treg homeostasis and fitness. Transplantation of wild type hematopoietic cells into a CD40-null recipient reduces Treg numbers allowing persistent T-cell activation and pro-inflammatory cytokines production causing, impaired B-lymphopoiesis. These evidences find correlation in aGVHD patients showing the loss of CD40+ BM-MSCs along with reduction in cells of the B-lineage. Modeling aGVHD in mice we show that the elimination of CD40+ BM-MSCs relies on their higher expression of MHC-I molecules. Indeed, aGVHD mice compared to MHC-matched controls showed the loss of MHC-I + radio-resistant host BM-MSCs. Our data point to CD40+ MHC-I+BM-MSCs as a key regulator of BM tolerogenic niches.

**Key points:**

1. CD40 regulates BM immunological tolerance following total body irradiation (TBI) and transplantation (BMT).
2. Loss of CD40^+^MHC-I^high^BM-MSCs is associated to BM manifestation of aGVHD in human and murine model.

## Introduction

Allogeneic bone marrow transplantation (BMT) remains the only therapeutic option for a wide range of hematological malignancies. However, it carries a significant risk of morbidity and mortality, which are mostly linked to adverse, immune-related events, such as infection and acute graft-versus-host disease (aGVHD). Thus, a better understanding of the mechanisms underlying immune regulation during bone marrow (BM) transplantation would significantly expand its application and potential benefits.

The BM has the features of an immune-privileged organ. This condition is granted by its peculiar enrichment in Foxp3+ regulatory T-cells (Tregs), which represent up to 30-40% of the total CD4 T-cell population, exceeding the fraction of Tregs in other lymphoid organs by 3- to 4-fold. In the BM, Tregs populate hematopoietic stem cell (HSC) niches where they protect endogenous HSCs from excessive inflammation and, during BMT, enable transplanted allogeneic HSCs to avoid rejection ^1 2^. Indeed, an excess of inflammatory cytokines, such as interferon (IFN) or tumor necrosis factor (TNF), can induce HSC dysfunction and hamper HSC differentiation into mature myelo-lymphoid lineages. After BMT, B-cells are particularly sensitive to the inflammatory conditions within the microenvironment and largely depend on the Tregs for proper reconstitution. In fact, B-cell lymphopoiesis depends on interleukin (IL)-7, which is defective in Treg-depleted mice ^3^. Furthermore, Treg depletion unleashes the production of granulocyte-macrophage colony-stimulating factor (GM-CSF), IL-6, and TNF, which promote granulopoiesis at the expense of B lymphopoiesis ^4^. Also IFN-γ negatively affects B-cell lymphopoiesis ^4,5^, suggesting that BM microenvironmental inflammatory state largely depend on Treg surveillance.

Tregs regulation has been largely studies in different tissues, but less so in the BM microenvironment. In the periphery, Treg development depends on transforming growth factor (TGF)-β and IL-2, while in the thymus IL-2 is dispensable because other coexisting cytokines can act on the IL-2Rβ chain ^6^. Further, the cluster of differentiation (CD)40/CD40 ligand (L) axis also contributes to the peripheral Treg pool ^7^.

Bone marrow mesenchymal stroma cells (BM-MSCs) nurse HSCs and progenitor cells through the stages of quiescence/self-renewal and differentiation ^8^. In addition to supporting HSC, BM-MSCs also exert immunoregulatory functions: they can inhibit the proliferation of T, B and natural killer (NK) lymphocytes ^9^ and promote the conversion of Teff to Treg, *in vitro*. On this line, *in vivo* administration of BM-MSCs has beneficial effects in the treatment of aGVHD ^10^. Beside the production of regulatory metabolic enzymes like indoleamine-2,3-dioxygenase (IDO) ^11^, the repertoire of co-stimulatory and/or co-inhibitory molecules harnessing BM-MSCs for immune regulation, including the role of CD40, is largely unknown. The lack of CD40 expression in normal BM and its upregulation on BM-MSCs of patients affected by splenic marginal zone lymphoma ^12^, suggest that BM-MSC can gain immunoregulatory properties in response to microenvironment perturbation.

In this study, we evaluated the role of CD40 on BM-MSCs in sustaining a BM tolerogenic state in the context of pro-inflammatory signals preventing unwanted T-cell activation that could negatively impact on hematopoiesis.

## Methods

### Patients

This study included bone marrow biopsies from 12 adult acute leukemia patients undergoing allo-HSCT at the Charité university hospital.

The patients’ clinical characteristics and GVHD prophylaxis are summarized in Table 1. The study was approved by the Charité-Berlin local ethics committee. Patients gave informed consent. The study was conducted in accordance with the Declaration of Helsinki”

### Mice

BALB/cAnNCrl and C57BL/6 mice were purchased from Charles River Laboratories (Calco, Italy). *Tnfrsf5* (*Cd40*)-KO mice (on a BALB/c background) were already available in our lab, while those on a C57BL/6 background were obtained from Dr. Dellabona Paolo (San Raffaele Hospital, Milan, Italy) and Prof. Bronte Vincenzo (University of Verona, Verona, Italy). CxB6F1 (*Cd40*)-KO mice were obtained by crossing the *Cd40*-KO on the BALB/c background with those on the C57BL/6 background. All experiments involving animals described in this study were approved by the Ministry of Health (authorization number n. 601/2019-PR).

### BMT and aGVHD mouse experiments

Canonical BMT experiments were performed by transplanting 2 × 10^5^ lin-cells into lethally irradiated WT and *Cd40*-KO mice as previously described ^13^. MHC-mismatched BMT was performed transplanting lethally irradiated BALB/c mice with 2 × 10^6^ T-cell-depleted BM cells from C57BL/6 mice in the presence or absence of Teff cells (2 × 10^5^) from the same donors. T-cell-depleted BM was obtained by flushing the BM cells from donors and incubating them with α-CD5 (Ly-1) microbeads from Miltenyi. Mice were assessed daily using a GVHD scoring system that measures items related to the known clinical signs of GVHD, including weight loss, posture, activity, fur texture, skin integrity, and diarrhea.

### Flow cytometry

The composition of the B-cell compartment in the BM and spleens of BM chimeras was analyzed as previously described ^14^. Notably, B-cell development in the BM has been classified into sequential subsets designated Fractions A, B, C, C’, D, E, and F as originally described by Hardy et al. ^15^.

Surface staining was performed in phosphate-buffered saline (PBS) supplemented with 2% fetal bovine serum (FBS) for 30 min on ice. Foxp3 intracellular staining was performed according to the manufacturer’s instructions (eBioscience). Before IFN-γ staining, the cells were stimulated *in vitro* for 4 h at 37°C with Cell Stimulation Cocktail plus protein transport inhibitors (eBioscience).

All antibodies are listed in Supplemental Table 2. Flow cytometry data were acquired on a LSRFortessa (Becton Dickinson) and analyzed with FlowJo software (version 8.8.6, Tree Star Inc.).

### Isolation and culture of murine BM-MSCs

Murine BM-MSC cultures were obtained from the trabecular fraction of femurs and tibias of WT and *CD40-*KO mice. Briefly, the cellular fraction of the femurs and tibias was washed out and the compact bone was incubated with collagenase I (1 mg/ml) for 1 h at 37°C. After enzyme digestion, the bone suspension was passed through a 70-mm filter mesh to remove any bone spicules and large tissues. Cells were seeded in complete medium at a density of 25 × 10^6^ cells/ml. Floating cells were removed every 3-4 days. Adherent cells were phenotypically characterized using the following antibodies: CD31, CD45, CD34, Ter119, CD44, Sca-1, and CD117. *In vitro* and *in vivo* experiments involving murine BM-MSCs were performed using cells between the 2nd and 5th passages.

### Statistical analysis

In transplantation analysis an ordinary one-way ANOVA was used, while for all the other comparisons the Unpaired t-test was used. Error bars in all graphs represent mean and standard deviation. All statistical analyses were performed with Prism 6 (GraphPad Software).

## Results

### Lethal irradiation perturbs BM T-cell homeostasis and promotes the expression of CD40 in stromal cells

We previously shown that CD40 expression characterizes stromal cells of patients who had the BM infiltrated by splenic marginal zone lymphoma. In these patient CD40 promotes a pro-inflammatory loop with mast cells, which sustain lymphoma cell growth ^12^. We analyzed the BM of mice developing autoimmunity and characterized for the expansion of IFN-γ + Teff ^16^. Stromal cells of these mice expressed CD40, compared to untreated mice (Online Supplementary Figure 1A). These data suggested that the presence of inflammatory cytokines such as IFN-γ could be relevant in increasing CD40 expression on BM-MSCs. Therefore we performed an *in vitro* analysis using primary human BM-MSCs and found that CD40 expression is induced by IFN-γ paralleling the induction of IDO1, a well know immune-regulatory molecules (Online Supplementary Figure 1B). These data suggested that MSC-derived CD40 could be relevant in the BM microenvironment when this microenvironment is perturbed toward the production of inflammatory cytokines.

**Figure 1.**
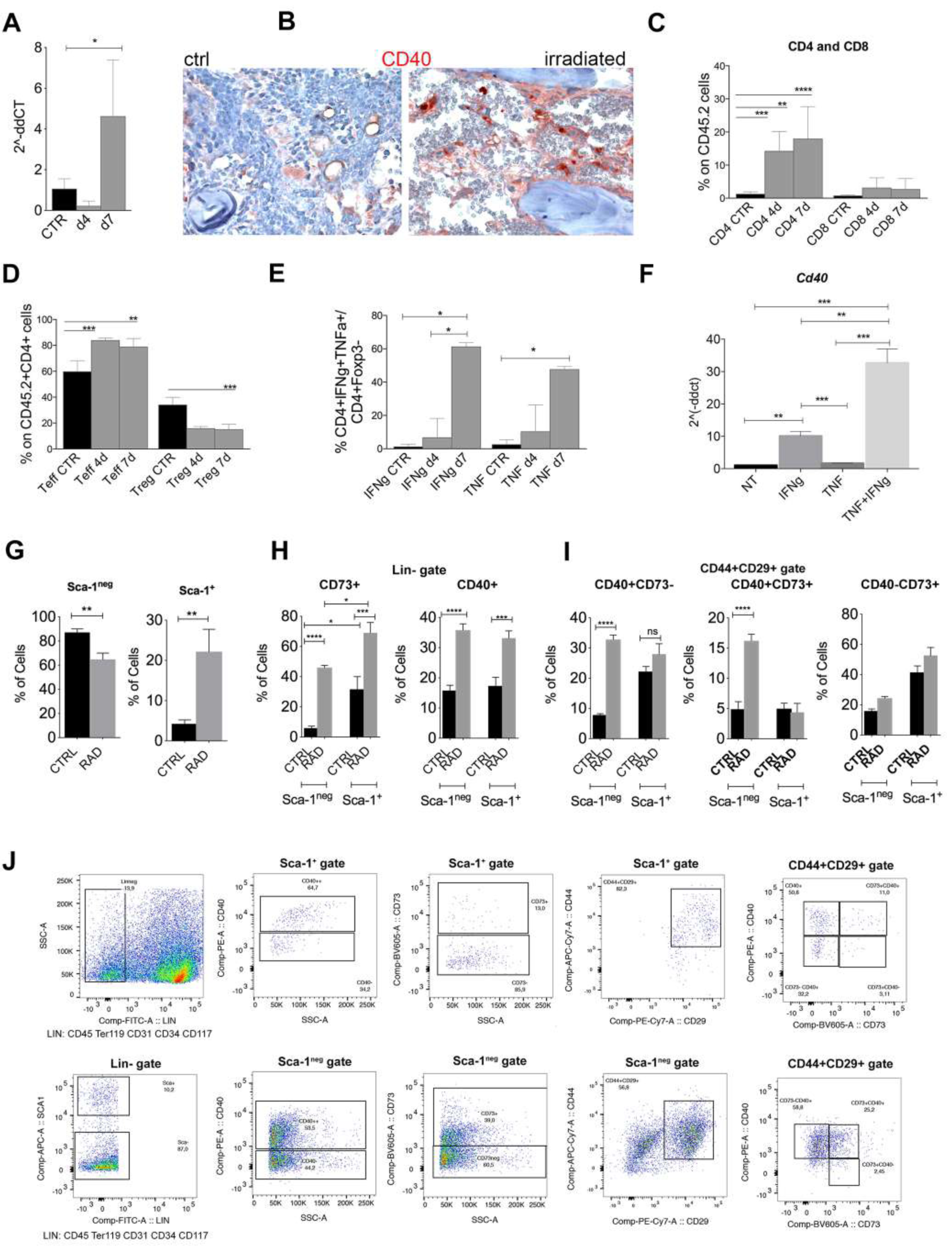
Expression of CD40 in the BM microenvironment. **A**. RT-PCR showing the expression of CD40 on primary BM-MSCs (not expanded *in vitro*) purified from BM at day 4 and 7 post-radiation and compared to basal. **B**. Representative image of CD40 IHC staining in the femurs and tibias of WT mice harvested 7 days post-irradiation. **C**. FACS analysis showing the frequency of CD4+ and CD8 T-cells in lethally irradiated (not reconstituted) mice (n = 5 per group). **p < 0.005, ***p < 0.001, compared by unpaired *t* test. **D**. Frequency of Teffs (CD25-Foxp3-) and Tregs (CD25+Foxp3+) in irradiated mice (CD4 gate) (n = 5 per group). **p < 0.005, ***p < 0.001, compared by unpaired *t* test. **E**. FACS analysis showing the frequency of Teff producing IFNγ and TNF (CD4+Foxp3-gate) (n = 5 per group). **p < 0.005, ***p < 0.001, compared by unpaired *t* test. **F**. Real-time PCR analysis for *Cd40* performed on total RNA isolated from *in vitro* expanded BM-MSCs stimulated 24 h with 10 ng/ml IFNγ and TNF or their combination (n = 5 per group). **p < 0.005, ***p < 0.001, compared by unpaired *t* test. **G**. Cumulative FACS analysis showing the frequency of Sca-1+ and Sca-1- BM-MSCs (CD29+CD44+CD45-Ter119-CD31-CD117-CD34- gate) in irradiated mice. **H**. Cumulative FACS analysis showing the frequency of CD73+ and CD40+ BM-MSCs within the Sca-1+ and Sca-1- gate. **I**. Cumulative FACS analysis showing the frequency of CD40+CD73-, CD40+CD73+ and CD40-CD73+ BM-MSCs in irradiated (RAD) vs non-irradiated mice (CTRL). N=8 for controls and n=10 for irradiated mice; **p < 0.005, ***p < 0.001, ****p < 0.0001, multiple comparisons using a one-way ANOVA. **J**. Representative gating strategy for CD73 positive, CD40 positive and CD40/CD73 double positive MSCs is shown.

Total body irradiation (TBI) is known to affect the BM immune microenvironment firstly inducing cytokines and then local immunosuppression to prevent the rejection of the donor bone marrow ^17^. We evaluated whether CD40 expression, undetectable in the BM-microenvironment in basal conditions, is induced by TBI. To this end BALB/c mice were lethally irradiated and sacrificed 4 or 7 days post-radiation to collect radiation-resistant BM-MSCs ^18^. Both RT-PCR (Figure 1A) on *ex-vivo* purified BM-MSCs (Online Supplementary Fig. 1C) and immunohistochemistry (IHC) (Figure 1B) showed that CD40 was overexpressed on total BM-MSCs at day 7 post-radiation compared to basal and day 4. The analysis of T-cell compartment showed the expansion of CD4+ cells upon irradiation both after 4 days and 7 days compared to non-irradiated mice (Figure 1C). Among CD4+ cells, we noticed that Teffs (CD4+Foxp3-) were increased in frequency compared to Tregs (CD4+Foxp3+) (Figure 1D). Moreover, the Teffs were activated to release TNF and IFNγ (Figure 1E), which, tested *in vitro*, were able to induce strong CD40 up-regulation in ex-vivo isolated and in vitro-expanded BM-MSCs (Figure 1F) in comparison to other stimuli also released in the BM of conditioned mice (i.e., G-CSF, GM-CSF+IL-6, and IL-17) ^19^ (Online Supplementary Figure 2A).

**Figure 2.**
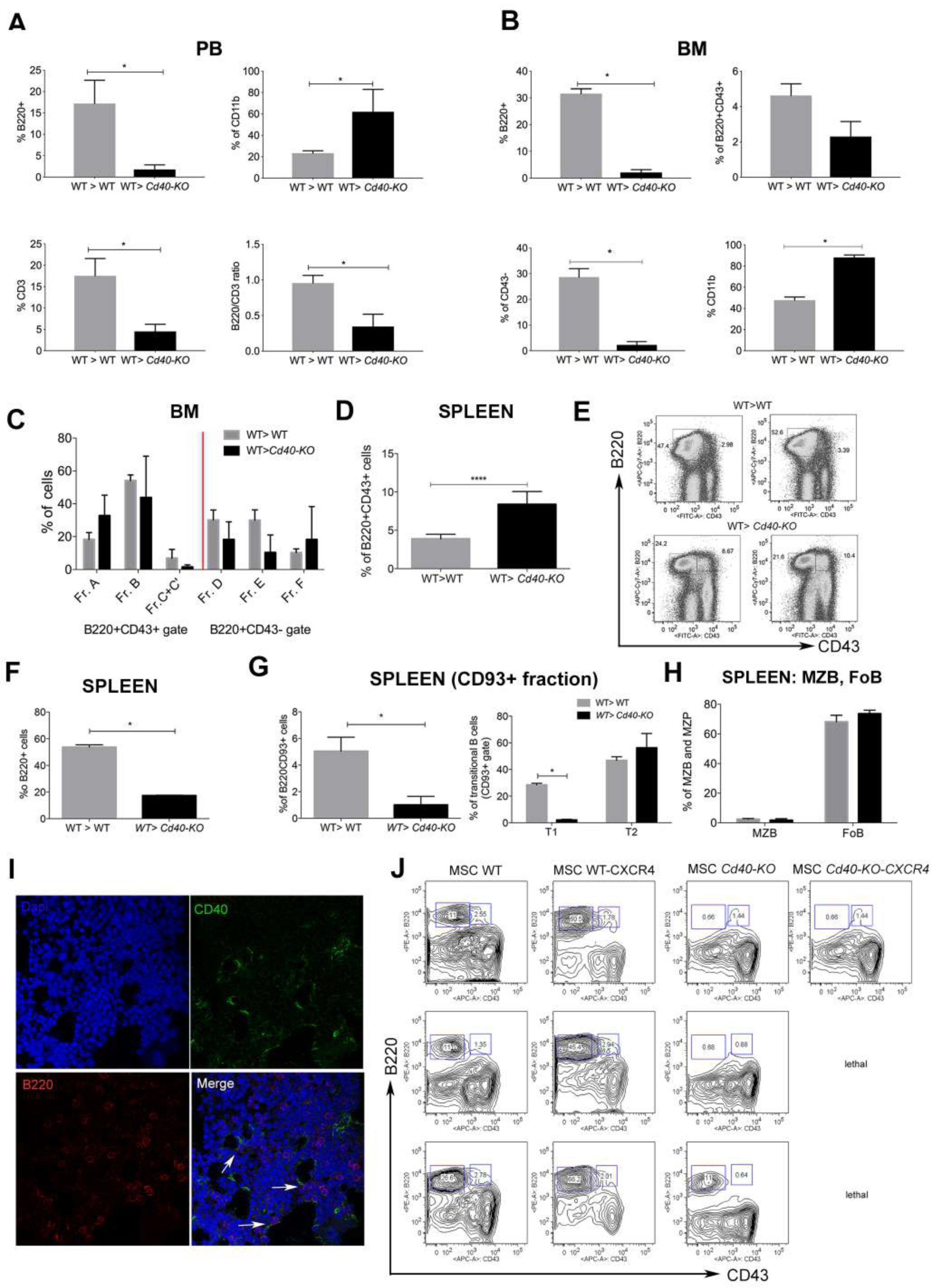
Analysis of B-cell development in WT>WT and WT>*Cd40*-KO BM chimeras. **A** Cumulative data for the PB FACS analysis showing the frequencies of B220+, CD3+, CD11b+ cells as well as the B220/CD3 ratio in the PB of WT>*Cd40-*KO compared to WT>WT BM chimeras. Representative plots and gating strategy are shown in Supplemental Figure 2a **B**. Cumulative data for FACS analysis of the BM showing the overall decrease in B220+CD43+ and B220+CD43 -B-cell subsets in *Cd40-*KO recipients along with an increase in the frequency of CD11b+ myeloid cells *p<0.05 **C**. Cumulative data showing the frequency of pre-pro-B and early pro-B precursors (A and B fractions), and of late pro-B and large pre-B precursors (C’-C) in chimeric mice; **D**. Cumulative data showing the frequency of B220+CD43+ B-cell precursors in the spleens of *Cd40-*KO recipient mice (n = 6 per group). **E**. Representative dot plots showing that B220+CD43+ cells are almost absent in the spleen of WT but not *Cd40-*KO recipients. **F**. Frequency of B220+ B-cells is lower in the spleen of chimeric mice than in WT mice. **G**. Frequency of B220+CD93+ and B220+CD93-B-cells in the spleen of chimeric mice. Splenic immature B220+CD93+ B-cell were divided into transitional T1, T2, and T3 cells based on their expression of CD23 and IgM (T1 = IgM + CD23-, T2 = IgM + CD23+, T3 = IgM^low^CD23+). **H**. Frequency of MZB and FoB within the gate of B220+CD93-cells is also highlighted. The relative gating strategy is shown in Supplemental Figure 2 *p < 0.05, compared using Student’s *t* test for all analyses. **I** Confocal microscopy analysis for CD40 (green) and B220 (red) expression showing the close contact between B220+ and CD40+elements in WT>WT BM chimeras. **J**. Dot plots showing B cell reconstitution in WT mice transplanted with Lin- cells co-infused with WT or *Cd40*-KO BM-MSCs transduced or not with CXCR4 (n=3/group).

To better characterize the BM-MSCs population expressing CD40 after TBI, we performed a multiparametric flow cytometry analysis. To define BM-MSCs we checked positivity for known surface MSC markers (CD44, CD29, Sca-1) and negativity for the lineage markers CD45, Ter119, CD34, CD117 and CD31. To our analysis we also added the CD73 marker, as it was recently attributed to radio-resistant and clonogenically-active BM-MSC ^20^. Our analysis showed that within the gate of CD29+CD44+CD45-Ter119-CD31-CD34-CD117-cells, lethal irradiation increased the frequency of Sca-1+ cells (Figure 1G). However, within both Sca-1+ and Sca-1-cells we found an increased frequency of CD73+ cells (Figure 1H). Paralleling CD73 we found that the frequency of CD40+ BM-MSCs was strongly increased after irradiation (Figure 1H). This suggests that CD40, as CD73, could be specifically induced and associated to radio-resistant BM-MSCs that could be either Sca-1+ or Sca-1-. Looking at the frequency of BM-MSCs co-expressing CD40 and CD73 within the gate of total BM-MSCs (CD29+CD44+CD45-Ter119-CD31-CD34-CD117-) we found an increased frequency of double positive CD40^+^CD73^-^ and CD40+CD73+ BM-MSCs upon irradiation (Figure 1I). The representative gating strategy is shown in Figure 1J.

To test whether the gain of immune regulatory features in BM-MSCs occured at the expense of their differentiation program toward osteo-and adipo-lineages, we performed RT-PCR analyses on TNF and IFN-γ stimulated cells. Osteoblast and adipocytes differentiation markers such as osterix, osteonectin, osteopontin, bglap, PPARg, were all down-modulated by TNF; osterix, Runx2, Sparc and Spp1 but not Bglap and PPARg were down-modulated by IFN-γ (Online Supplementary 2B). The combination of TNF and IFN-γ was additive in decreasing the expression of *Spp1* (Online Supplementary 2B). As control, the same MSCs showed increased CD40 expression, suggesting an inverse relationship between the expression of immune and differentiation programs.

### Altered B-cell lymphopoiesis in the BM is a characteristic of Cd40-KO recipient chimeric mice

To study the regulatory role of CD40 on BM-MSCs *in vivo*, we set up bone marrow transplant (BMT) experiments in which recipient mice (either *Cd40-*KO or WT, all CD90.2) were lethally irradiated and transplanted with HSCs from congeneic CD90.1 WT donors. Twenty-one days post-BMT we performed FACS on peripheral blood (PB), BM, and spleens of recipient mice. Compared to WT, *Cd40-*KO recipients showed a prominent decrease in the frequency of PB B-cells and, to a lesser extent, of T-cells, but also an increased frequency of CD11b+ myeloid cells (Fig. 2A, Online Supplementary 3A). In line with our PB analysis, we observed a reduction of lymphoid cells in the BM of *Cd40-*KO recipients despite normal myelopoietic (CD11b cells) development (Figure 2B). Particularly, B-cell development was largely defective, as shown by the reduced B220+CD43+ and almost absent B220+CD43-fractions (Figure 2B). B-cell development was arrested at the A and B fractions, which correspond to the pre-pro-B and early pro-B phases, respectively, with reduced development of the fraction of C’-C precursors (late pro-B and large pre-B) (Figure 2C; Online Supplementary 3B) ^15^. Intriguingly, in the spleens of *Cd40-*KO recipients we noted an increased frequency of B220+CD43+ B-cell precursors that developed into mature B-cells of different types, thus giving rise to extramedullary B lymphopoiesis (Figure 2 D-E). However, the overall frequency of B220+ cells was significantly reduced in the spleen of *Cd40*-KO recipients (Figure 2F) with a significant impairment in the CD93+ immature fraction (particularly affected were transitional T1 cells, Figure 2G). Nevertheless, the presence of mature marginal zone B-cell and follicular B-cell (Figure 2H; Supplemental Figure 3C) suggested a compensatory role for splenic B-cell lymphopoiesis in response to dysfunctional BM B-lymphopoiesis. Notably, the defective B-cell lymphopoiesis in the *Cd40-*KO recipient chimeras was not due to a systemic anti-CD40 response as CD40+ B-cells were still present in the spleen and PB (Online Supplementary Figure 4A). The relevance of BM-MSC-derived CD40 in B cell development was also suggested by the close contact between CD40+ mesenchymal cells and B220+ cells *in vivo* as detected by confocal microscopy analysis for B220 and CD40 on BM biopsies (Figure 2I). We functionally proved this hypothesis by performing a BMT experiment where WT mice were lethally irradiated and reconstituted with Lin-cells in the presence of either WT or *Cd40-*KO BM-MSCs. BM-MSCs were also forced to express CXCR4 to promote BM homing. The presence of WT CXCR4+ BM-MSCs promoted a higher frequency of B220+ cells compared to mice that did not received MSCs (Figure 2J) at day 14 post-BMT. On the contrary, *Cd40*-KO BM-MSC negatively affected B cell development and 33% of mice died (Figure 2J).

**Figure 3.**
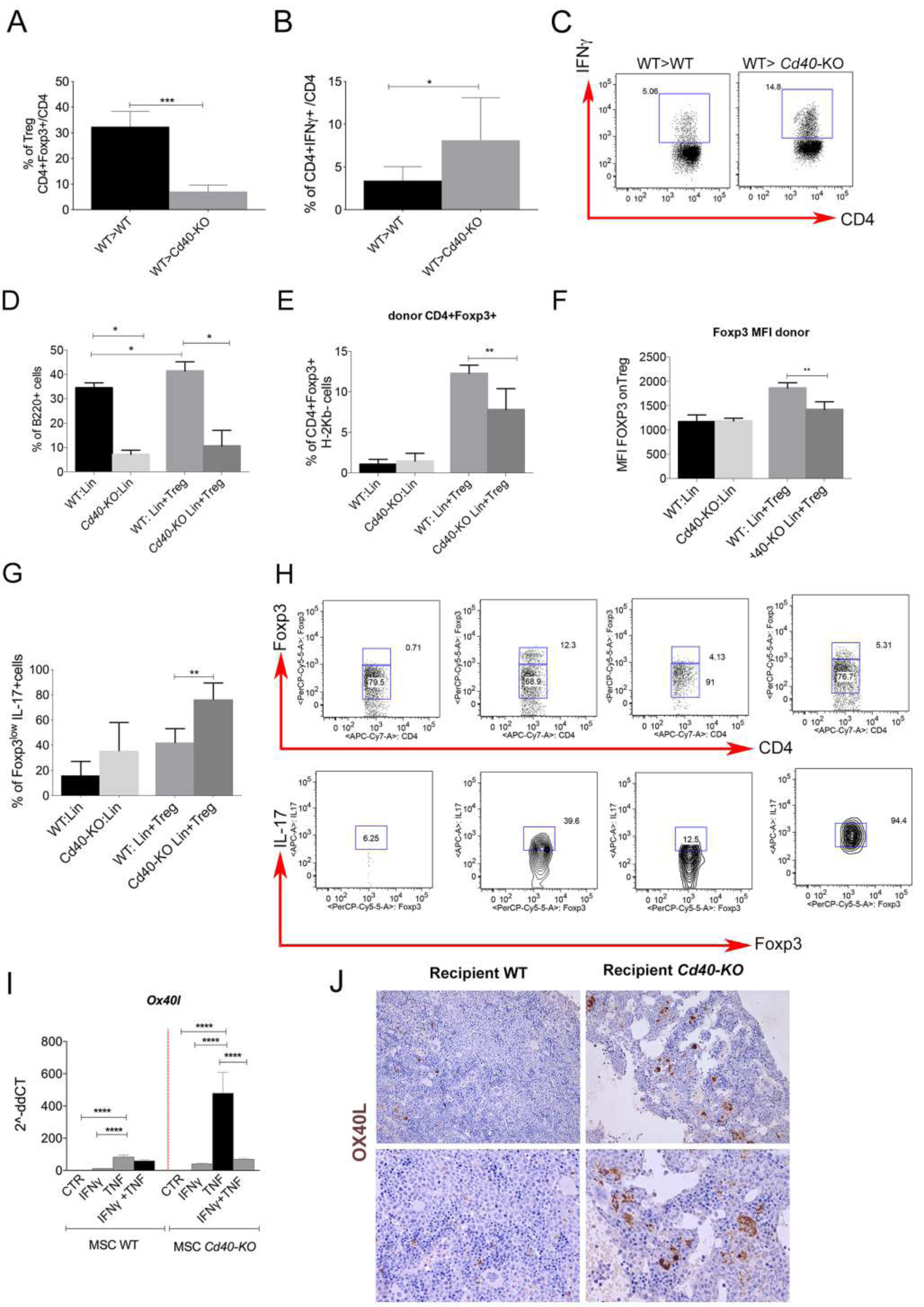
Characterization of T-cell status in the *Cd40*-KO recipient BM chimeras. A. Treg frequency in the BM of WT>WT and WT>*Cd40-*KO BM chimeras (n = 6 per group). ***p < 0.001, compared using Student’s *t* test. **B**. Cumulative data and **C**. representative dot gating strategy showing the production of IFNγ by CD4+Foxp3-Teffs s in the BM of WT>WT and WT>*Cd40-*KO BM chimeras (n = 6 per group). *p < 0.05, compared by Student’s *t* test. **D**. Frequency of B220+ cells in the WT>WT and WT>*Cd40-*KO BM chimeras after lethally irradiating CxB6F1 WT or *Cd40-*KO mice with lin-cells and spleen-derived Tregs from donor BALB/c mice (n = 5 per group). *p < 0.05, compared by Student’s *t* test. Please see Supplemental Figure 4a for a schematic representation of this experiment. **E**. Frequency of donor (H-2Kd^+^Kb^-^) CD4+Foxp3+ Tregs in the WT>WT and WT>*Cd40-*KO BM chimeras reconstituted with lin- and Tregs (n = 5 per group). *p < 0.05, compared by Student’s *t* test. **F**. Cumulative data showing the MFI of Foxp3 on Treg cells. **G**. IL-17 production by Foxp3^low^Tregs in *Cd40-*KO recipients (n = 5 per group). **p < 0.005, compared by Student’s t test. **H**. Representative gating strategy for Foxp3 and IL-17 analysis in the different BM chimeras. **I**. Quantitative RT-PCR analysis of *Ox40l* gene expression in WT and *Cd40-*KO BM-MSCs treated *in vitro* with IFN-γ, TNF, or their combination (n = 5 per group). ****p < 0.0001, multiple comparisons using a one-way ANOVA. **J**. Representative images of OX40L staining in BM sections of the WT>WT and WT>*Cd40-* KO BM chimeras. Mesenchymal-like elements (arrows) are shown in greater detail in Supplemental Fig. 4c.

**Figure 4.**
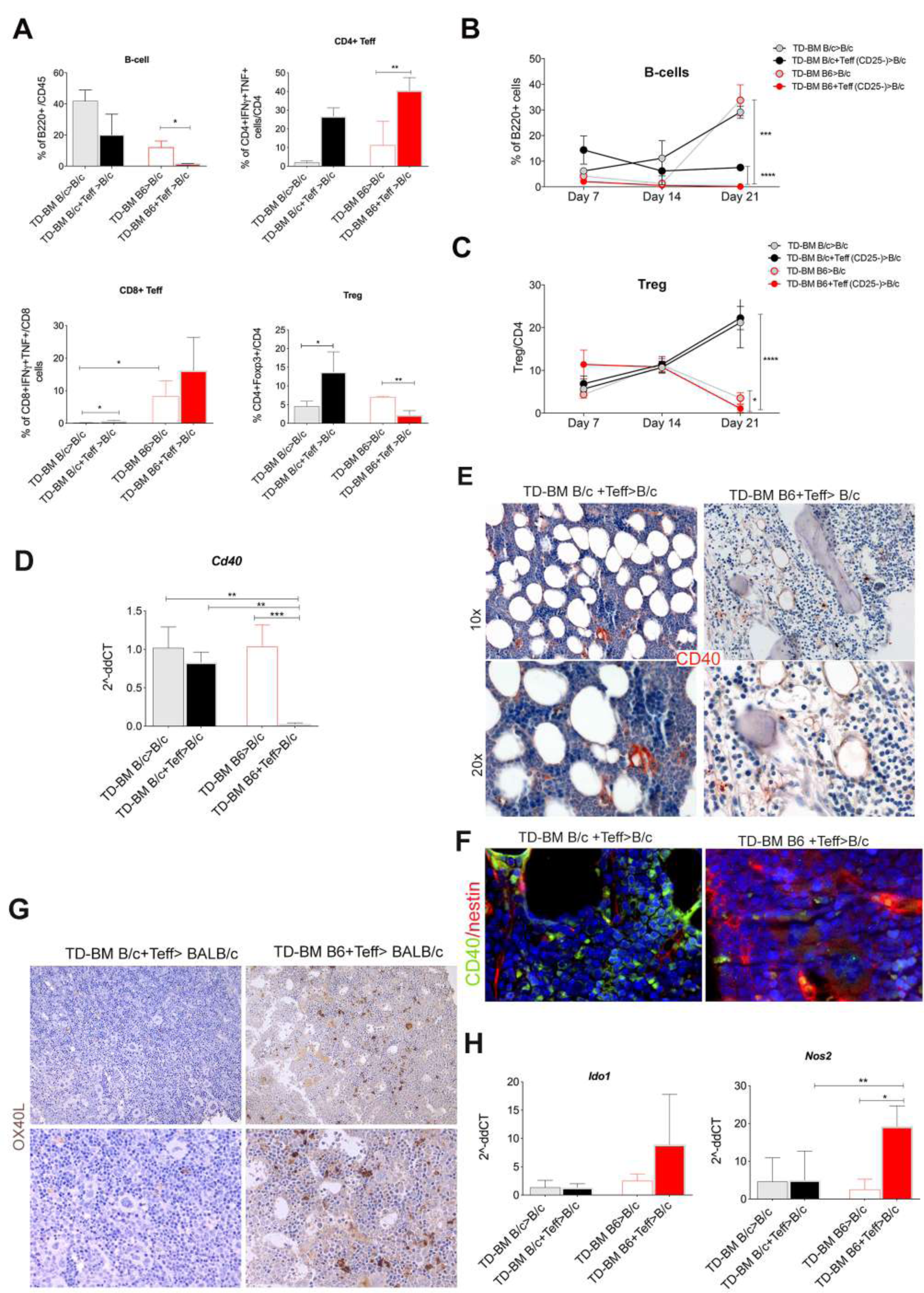
Downregulation of CD40 expression in the BM-MSCs of aGVHD mice. **A**. Frequency of B220+ B-cells, CD4+Foxp3- Teffs releasing IFNγ and TNF, CD8+ T-cells releasing IFNγ and TNF and CD4+Foxp3+ Tregs in the aGVHD mice compared to control animals. *p < 0.05, **p < 0.005, ***p < 0.001, compared by Student’s *t* test. PB FACS analysis showing changes in the frequency of **B**. B-cells and **C**. Tregs at days 7, 14, and 21 post-allogeneic BMT compared to controls (n = 5 per group). ***p < 0.001, compared by Student’s *t* test. **D**. qPCR analysis of *Cd40* expression in BM-MSCs isolated from aGVHD and control mice. **p < 0.005, ***p < 0.001, compared by Student’s *t* test. **E**. Representative CD40 IHC staining in aGVHD mice (TD-BM B6 + Teff>BALB/c) compared to controls (TD-BM BALB/c + Teff>BALB/c). **F**. Representative co-immunofluorescence showing CD40 (green) and nestin (red) levels in aGVHD mice (TD-BM B6 + Teff>BALB/c) compared to controls (TD-BM BALB/c + Teff>BALB/c). **G**. Representative images of OX40L IHC staining in BM sections from allogeneic-transplanted animals (TD-BM + Teff B6>B/c) or controls (TD-BM BALB/c + Teff>BALB/c). **H**. Quantitative RT-PCR analysis of *Ido1*, and *Nos2* expression in BM-MSCs isolated from allogeneic-transplanted animals (TD-BM + Teff B6>B/c) and controls (TD-BM BALB/c + Teff>BALB/c).

### CD40-deficiency in the BM stromal compartment creates a lymphopoietic niche unfit to respond under stress condition

We next tested whether the lack of stromal CD40 expression might negatively impact on Treg development or generate a pro-inflammatory microenvironment, which in turn can inhibits B-cell development.

To this end, we analyzed BM T-cell status in WT> *Cd40-*KO and WT>WT chimeras, 4 weeks after BMT. Tregs were reduced in the BM of WT> *Cd40-*KO chimeras compared to the WT>WT counterpart (Figure 3A), and IFNγ-producing Teff cells were significantly increased (Figure 3B, C). To gain further insight into the behavior of Tregs in the *Cd40-*KO recipients and their relationships with B-cell development, Tregs were co-infused with donor lin-cells (both B/c, H-2d) into either WT or *Cd40-*KO F1 recipients (both CxB6, H-2b/d) (Online Supplementary Figure 5A). Of note, Tregs accelerated the recovery of B-cells in BM of WT>WT chimeras (Figure 3D), but not in *Cd40-*KO recipients. In line with this result, a reduced frequency of donor Tregs was observed in *Cd40-*KO compared to WT recipients (Figure 3E). Such reduction of donor Tregs was paralleled by decreased Foxp3 levels (Figure 3F) and increased production of IL-17 in the CD4^+^Foxp3^low^ population (Figure 3G, H) in *Cd40-*KO recipients. This phenotype suggests a possible conversion of Tregs into Th17 cells ^21^. In line, transplantation of WT donor cells into *Cd40-*KO recipients resulted in reduced OX40 expression on Tregs, a known marker of fitness ^22^ (Online Supplementary Figure 5B). To try to explain this phenotype, we evaluated OX40L expression in *Cd40-*KO and wt BM-MSCs. Indeed, previous data have shown that in the absence of an intact CD40/CD40L axis, OX40 triggering could worsen allograft rejection in cardiac transplant model, through the exacerbation of the Th1 and Th2 responses ^23^. RT-PCR analysis performed onto *ex-vivo* stimulated BM-MSCs showed that *Cd40-*KO BM-MSCs had higher OX40L levels than their WT counterparts in response to TNF (Figure 3I). In line, BM IHC for OX40L showed an overall increase in its expression in *Cd40-*KO recipient mice (Figure 3J), and confocal microscopy analysis revealed co-staining with the nestin marker (Online Supplementary Figure 5C). These data could explain the observed impairment of Tregs in favor of Teffs in the WT>*Cd40-*KO chimeras. In fact, the increased OX40L stimulation was previously shown to promote the conversion of Tregs to Th17 ^24^. This is compatible with our earlier findings of increased production of IL-17 in Foxp3^low^ cells of *Cd40-*KO recipient mice. Conversely, upon OX40L stimulation, Teff cells increase proliferation and acquire memory functions ^25,26^. Overall, these data suggest that in the absence of CD40, BM-MSCs are unable to regulate donor Treg activity. Indeed, under certain inflammatory stimuli Treg may might lose FOXP3 expression and acquire the ability to produce Th cytokines ^27^.

**Figure 5.**
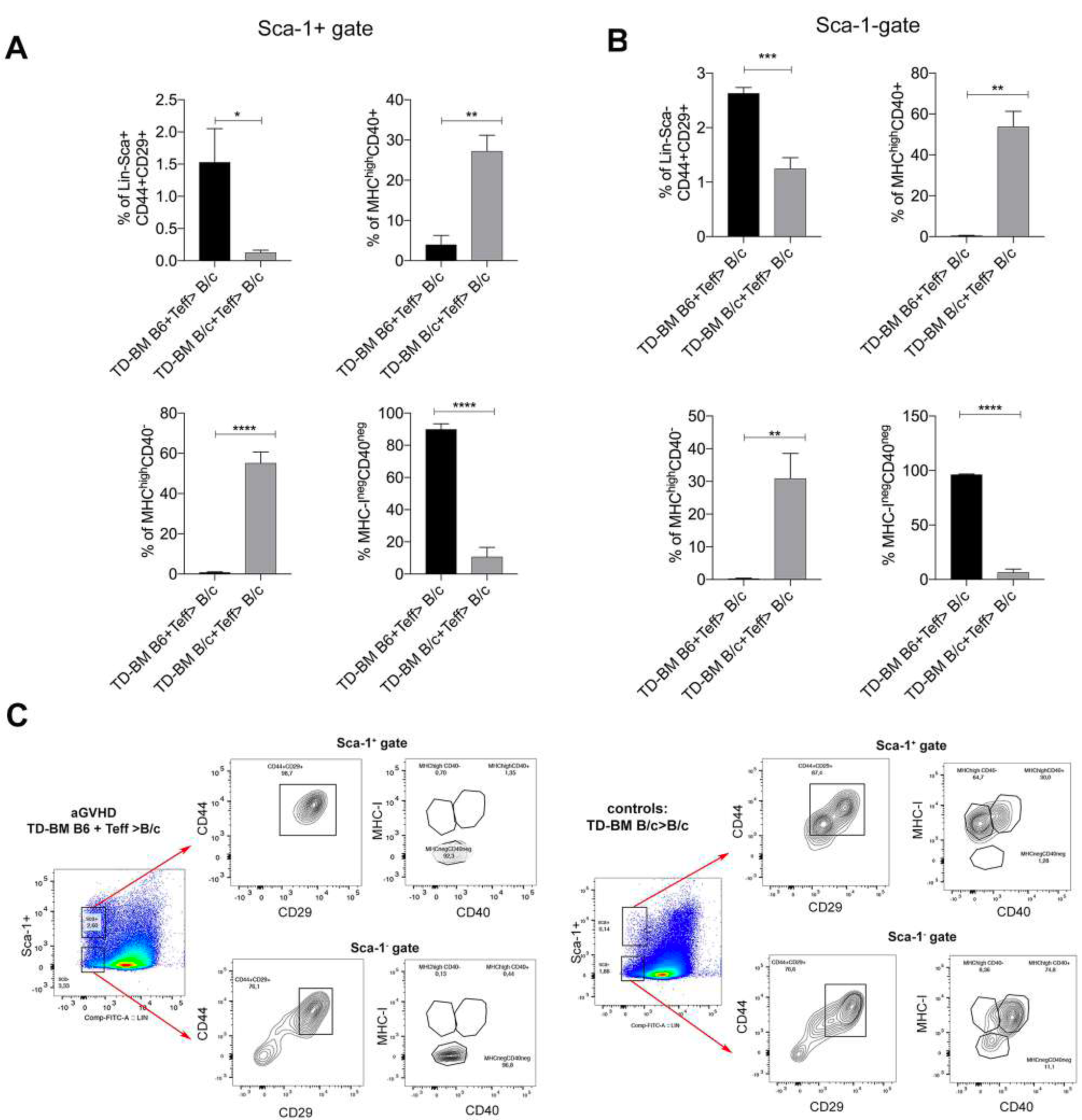
MHC-I^high^ MSCs are specifically eliminated in aGVHD mice. **A**. Frequency of MHC^high^CD40+, MHC^high^CD40- and of MHC^neg^CD40+ BM-MSCs in the Sca-1+ gate and **B**. Sca-1^neg^ gate of Lin-CD44+CD29+ BM-MSCs in aGVHD vs controls mice. The analysis was performed at 14 days post-transplantation. *p < 0.05, **p < 0.005, ***p < 0.001, compared by Student’s *t* test, ****p < 0.0001, compared by Student’s *t* test. **C**. Representative gating strategy for the above analysis.

### Stromal CD40 expression is downregulated during BM manifestation of acute-GVHD disease

Allogeneic HSC transplantation is an effective treatment for many hematologic diseases, but one major treatment-associated complication is aGVHD. aGVHD patients often show impaired B-cell immunity as well as activated donor T-cell-mediated osteoblast destruction ^28 29^. Given the striking similarity of this phenotype with our findings in the BM of irradiated *Cd40*-KO mice, we set up experiments to evaluate whether the loss of CD40 expression on BM-MSCs could be a common feature in BM experiencing an abnormal T-cell activation, such as during aGVHD pathogenesis. To model GVHD, we used an experimental model in which major histocompatibility complex (MHC)-mismatched BM cells and lymphocytes isolated from C57BL/6 (H2^b^) donors were transplanted into BALB/c (H2^d^) recipients. 2 × 10^6^ T-depleted donor BM (TD-BM) cells were co-infused or not (control) with 5 × 10^5^ donor CD4+CD25- (Teff) lymphocytes ^30^. As a further control, a non-MHC mismatched BMT was also performed using BALB/c (H2^d^) mice as both donor and recipient (Supplemental Fig. 6A). Under the allogeneic conditions, aGVHD manifested within 21 days as confirmed by changes in mice weight (Online Supplementary Figure 6B) and histological analysis of the liver, skin and lung, in accordance with a murine grading system (Online Supplementary Figure 6C-D) ^31^. FACS analysis of BM cell suspensions isolated from these mice revealed impaired B-cell development associated with an increased production of IFN-γ and TNFα by CD4 and CD8+ cells (Figure 4A). Notably, Treg frequency was significantly reduced in mice with aGVHD. Similar changes in the B-cell and Treg populations were also detected in PB samples, where a time-dependent loss of both cell types was observed (Figure 4B-C). These data suggest that Treg and B-cell frequencies could be used as early markers to identify patients who will likely develop aGVHD.

**Figure 6.**
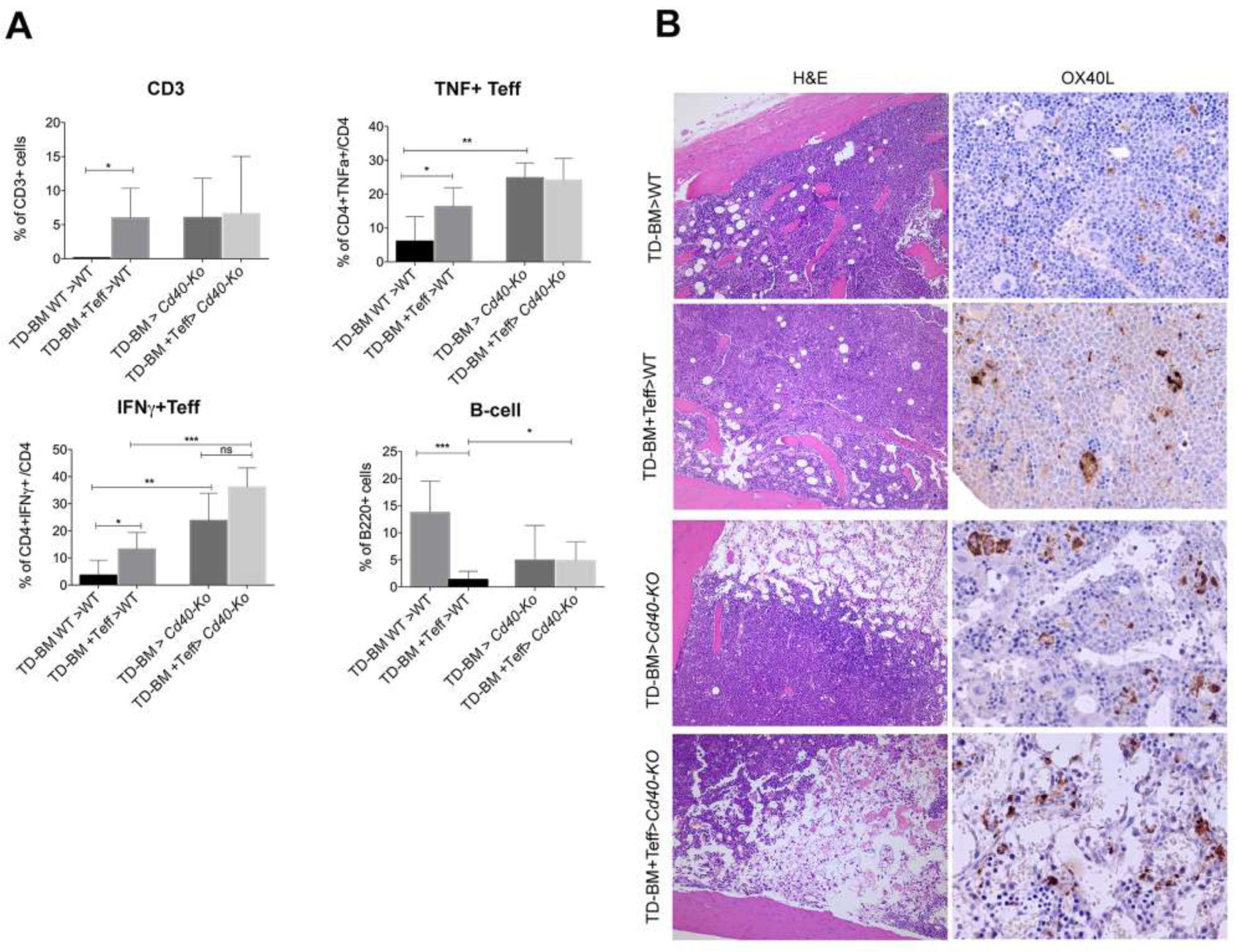
OX40L expression is increased in the BM of aGVHD mice. **A**. Frequency of CD3, IFNγ+ and TNF+ Teff and B220+ cells in *Cd40-KO* and WT mice receiving an MHC-mismatched BM. *p < 0.05, **p < 0.005, ***p < 0.001, compared by Student’s *t* test. **B**. Representative H&E and OX40L IHC images of BM sections from aGVHD (TD-BM + Teff B6>B/c) *vs* controls (TD-BM BALB/c + Teff>BALB/c) chimeras.

We next analyzed CD40 expression in the BM microenvironment during aGVHD. RT-PCR analysis showed loss of CD40 expression in freshly isolated BM-MSCs from aGVHD but not control chimeras (Figure 4D). IHC of the BM specimens confirmed CD40 loss during aGVHD (Figure 4E) and co-immunofluorescence analysis for nestin and CD40 confirmed the lack of CD40 on nestin+ stromal cells from aGVHD mice (Figure 4F). Notably, the lack of CD40 was coupled with increased OX40L on BM-MSCs (Figure 4G), which also showed increased production of NOS2 and IDO-I (Figure 4H). These data suggest that aGVHD development is associated with an altered immunoregulatory axis involving CD40 expression on BM-MSCs and Treg. The increased expression of *Ido1* and *Nos2* in BM-MSCs from aGVHD mice suggests that different mechanisms are in place to try to restore immune homeostasis. However, despite these mechanisms the outcome of BMT in CD40-deficient mice suggests that they cannot bypass the requirement for a functional CD40-CD40L axis.

### Loss of MHC-I^high^ MSCs in BM from aGVHD mice

To try to explain the loss of CD40+ BM-MSCs we evaluated the expression of MHC-I, which is necessary for allo-recognition on BM-MSCs from lethally irradiated and aGVHD (TD-BM B6+Teff>B/c) *vs* controls (TD-BM B/c+Teff>B/c) mice. Lethal irradiation increased the expression of MHC-I in Sca-1-CD29+CD44+ BM-MSCs (Online Supplementary Figure 7). A similar trend was observed in Sca-1+ cells. Both Sca-1+ and Sca-1^-^ comprised CD40+ elements (not shown). The same analysis performed onto aGVHD *vs* controls mice, at 14 days post-transplantation, revealed a dramatic change in term of MHC-I+ cells composition in both the Sca-1+ and Sca-1-fraction. Indeed, in both populations we found an almost complete loss of MHC-I^high^CD40+, and MHC^high^CD40-subsets, notably leaving intact those expressing lower level of MHC-I (Figure 5A-C).

**Figure 7.**
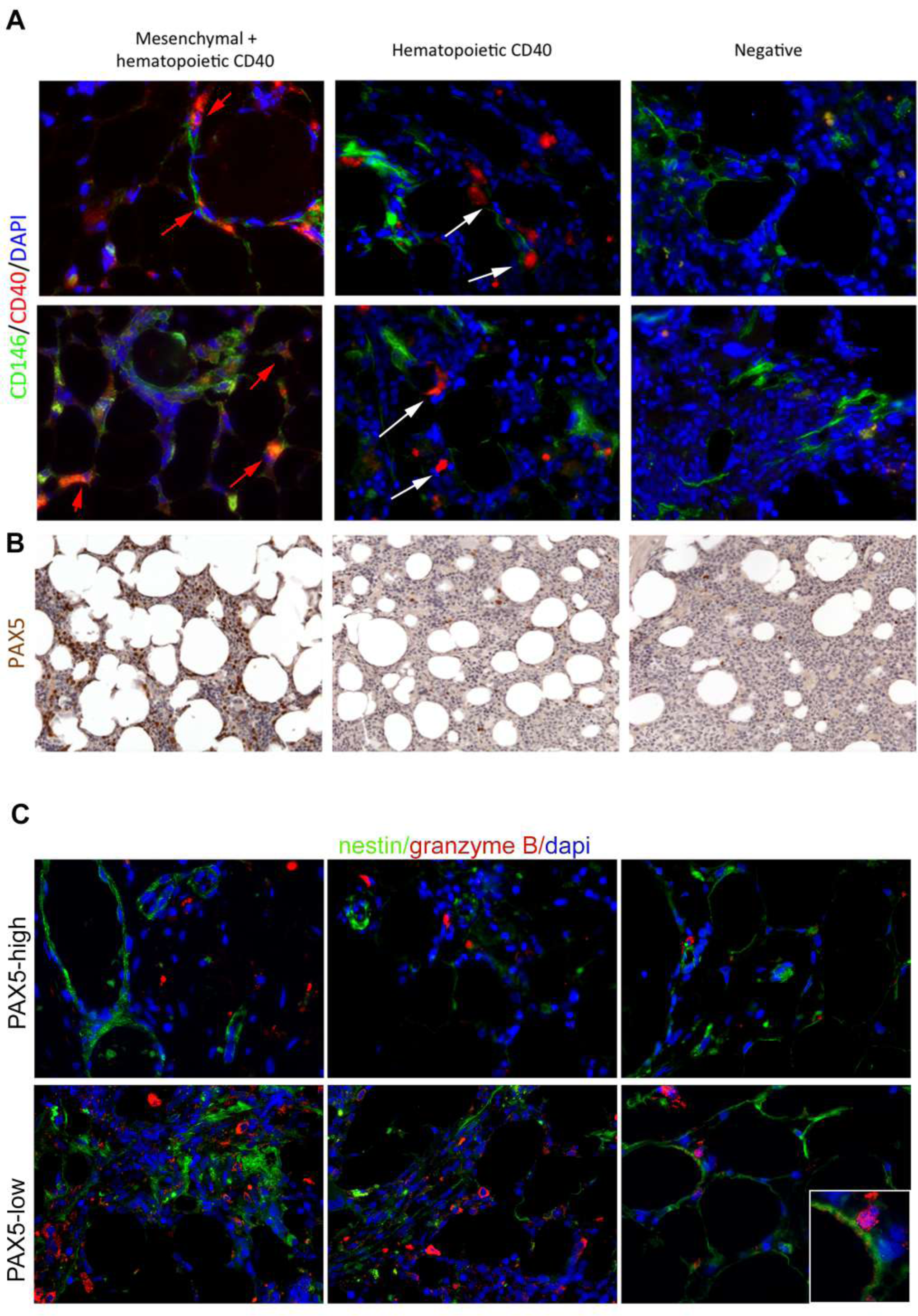
Lack of CD40+ BM-MSCs in BOM from aGVHD patients. **A**. Representative double-marker immunofluorescence analysis for CD40 (red) and CD146 (green), showing CD40 expression shared by CD146+ mesenchymal elements (red arrows) or confined to CD146-hematopoietic cells (white arrows). **B**. Pax5 IHC showing a variable expression of Pax5 with the highest fractions of Pax5+ cells observed in cases in which CD40 was expressed also in the mesenchymal cells. Patient’s characteristics and quantitative IHC data are included in Supplemental Table 1.

### The absence of CD40 specifically worsen the BM manifestation of aGVHD

To assess the relevance of CD40 in the context of aGVHD, *Cd40-*KO and WT mice were transplanted with MHC-mismatched TD-BM cells co-infused or not with donor CD4+CD25-Teff. We found that transplantation of *Cd40-*KO mice with a MHC-mismatched BM was sufficient to promote an excessive production of inflammatory cytokines in the BM even in the absence of donor Teff (Figure 6A), so much that recipient mice were sacrificed at day 14 instead of day 21 because of their suffering conditions (at day 12, 15 % of *Cd40-*KO recipients were died). In these mice, FACS analysis of BM cells showed a significant increase in the frequency of CD3+ T cells indicating that the *Cd40-*KO microenvironment favors the proliferation and pro-inflammatory activation of radioresistant host T cells (but also donor T cells, when co-infused). Indeed, we found high production of IFN-γ and TNF by CD4-T cells along with decreased frequency of B220+ cells (Figure 6A). Notably, the reduced frequency of donor TD-BM cells in *Cd40-*KO compared to WT recipients (not shown), suggests that the *Cd40-KO* microenvironment might favour the generation of a graft vs donor response.

Histological and IHC analysis showed a reduced BM cellularity and increased expression of OX40L on mesenchymal elements of *Cd40-*KO recipients (Figure 6B). Notably, the exacerbated inflammatory condition due to CD40 deficiency in radio-resistant cells, both in presence and absence of donor Teff, was restricted to the BM environment, as the histology of infiltrating lymphocytes in peripheral organs (liver and skin) was not different between WT and *Cd40-*KO recipients (Online Supplementary Figure 8).

### Association between CD40 and PAX5 loss in aGvHD patients BOM

To assess the relevance of our findings in human disease, we examined BM biopsies from 12 allo-HSCT patients, seven of which developed aGVHD (grade I-III). Patient’s characteristics are included in Supplemental Table 1. BM Paraffin sections were stained with Ab to CD40 and to PAX5, an early B-cell transcription factor expressed since the early stages of B-cell lymphopoiesis. A blinded analysis of these cases showed that CD40 was either not expressed, or expressed on scattered elements with myeloid morphology, or in spindle- or stellate-shaped stromal cells in other different cases. The mesenchymal nature of the spindle- or stellate-shaped cells was confirmed by double-marker immunofluorescence analysis for CD40 and CD146 (Figure 7A). Pax5 expression ranged from less than 1% to nearly 20% of the hematopoietic cellularity and positively correlated with CD40 expression on mesenchymal cells. B-cell frequencies were lower in cases of non-stromal or negative expression of CD40. The last condition characterized GVHD patients (Figure 7B and Online Supplementary Table 1) in full agreement with the described mouse data. The analysis of the expression and localization of granzyme-B in BOM from GVHD patients showed that patients that had a strong reduction in PAX5+ cells had granzyme B+ granules localized onto CD146+ BM-MSCs cells (Figure 7C), suggesting that Teff cells are actively killing BM-MSCs.

## Discussion

The bone marrow (BM) is a peculiar lymphoid organ. Mature and memory T cells relocate in the BM parenchyma, which is a potential source of effector T cells ^32^. This could, at least in part, explain why the BM is the organ with the highest frequency of Treg compared to other secondary lymphoid organs, such as spleen and LN ^33^. Furthermore, the strong detrimental effect of uncontrolled cytokine production over HSC homeostasis and differentiation might explain the need of high Treg activity to keep effector T cells silent ^1^. This is particularly evident for B-cells. In fact, the initial phases of B-cell development is influenced by the BM immune environment. In fact an excessive IFN-γ production ^5^ or changes in Treg composition ^3^ has been shown to negatively affect B-cell development.

Here, we provide evidence that the immune regulatory activity of BM-MSCs over Treg and Teff, which is generally ascribed to soluble molecules (i.e. IDO, NOS-2), relies on the expression of CD40, a cell surface receptor belonging to the TNF-R family which function is critical for the antigen presenting cells activity ^34^. Starting from our previous finding of CD40 up-regulation in CD146+ mesenchymal cells of SMZL patients, we show here that CD40 expression in the BM microenvironment is induced under inflammatory conditions, such as those associated with lethal irradiation. Modelling BMT in mice, we show that the post-radiation inflammatory response, guided by IFN-γ and TNF, is sufficient to induce CD40. BM-MSCs and Foxp3+ Treg cells closely interact to confer immune privilege to BM ^35^. We showed that the prompt up-regulation of CD40 in radio-resistant BM-MSCs during TBI, is required to grant Treg functions. Accordingly, the absence of CD40 on stromal elements reduces Tregs in favor of Teff, possibly via conversion to Th17 cells.

This condition was associated to defective B-cell development in *Cd40*-KO mice, mirroring the prolonged B-cell dysfunction, the increased T-cell infiltration and the disrupted BM niche that characterize human aGVHD ^29.^ Indeed, BM biopsies from aGVHD patients reveal lack of PAX5 expression, a marker that identifies early B-cells, along with the loss of stromal CD40 expression.

Mechanistically, we found that the absence of CD40 on stromal cells was associated to higher level of OX40L expression, a feature underestimated in BM-MSCs. Our data suggest that in the absence of CD40/CD40L, the OX40/OX40L axis might prevail. In line, stimulation of OX40 overrides cardiac allograft acceptance induced by disrupting CD40-CD40L interaction ^23^. In the BM this might favor Teff proliferation at the expense of Treg, thus creating a pro-inflammatory environment that in case of MHC-mismatch, might favor the development of aGVHD.

The loss of CD40+ BM-MSCs in aGVHD, might be due to their high MHC-I expression. Indeed, modeling GVHD in mice we found a time-dependent loss of MHC-I-high MSC subset, which comprises CD40+ cells. This finding opens two possible scenarios: the first one is the existence of a MHC-driven allo-recognition and therefore elimination of MHC-I^+^ BM-MSCs, and the second one is the possible down-modulation of MHC-I molecules on BM-MSCs, to prevent allo-recognition. BM-biopsies from aGVHD patients showing the disruption of osteoblasts of the BM niche ^29^ together with our data showing the co-localization of granzyme-B granules with nestin+ BM-MSCs in the same aGVHD patients strongly support the first hypothesis.

Furthermore, analyzing the compartment of radio-resistant BM-MSCs we show that lethal irradiation increases the frequency of CD40+ BM-MSCs and that a subset of these cells are CD73+. Indeed, CD73 is an ectoenzyme involved in the production of adenosine, a potent immune suppressive molecule, also affecting Treg ^36^. This might explain why the loss of CD40+ BM-MSCs subset during GVHD results in exacerbated inflammatory response.

Different clinical trials testing the efficacy of BM-MSCs to prevent or treat GVHD are running ^37^. Most of them utilize allo-MSCs from third-party donors. Results from these trials show no collateral side effects and a good overall response rate in chronic GVHD patients. In aGVHD patients no significant differences have been shown among MSCs vs no MSCs groups, although encouraging results have been reported in one controlled clinical trial with 40 patients in which a complete or partial response was observed in 15 and 1 patient, respectively. Several clinical trials are still on going and will provide additional data to evaluate the efficacy of allo-MSC administration. Nevertheless, some key factors that can influence MSCs activity have been identified, such as the route of administration and the source of allo-MSCs. On this regards our data provide new insight suggesting that although the total MSCs population is potentially able to provide immune suppression, only the specific subset of BM-MSCs expressing CD40 could be the most effective for the generation of BM tolerance. However, their tolerogenic effect could be reduced because of their high expression of MHC-I molecules, which may lead to their elimination.

Finally, the study shows that the gaining of regulatory functions along with CD40 upregulation, in MSCs is associated with the down-regulation of their speciation markers (Bglap, Osterix, Osteopontin) in favor of immune regulatory feauture. This is either explained by a more undifferentiated/progenitor subset of BM-MSCs exerting these functions, or by a switch from an architectural/supportive hematopoietic niche function of BM-MSCs to an immunoregulatory activity when excessive inflammatory conditions are sensed. Additionally, here we described that high expression of CD40 is on Sca-1+, a marker that identifies stromal progenitors able to generate both osteogenic and stromal cells and able to provide a supportive environment for hematopoiesis ^38^. It has been reported that Sca-1-positive cells have higher colony forming ability and show enhanced proliferation compared with Sca-1-negative cells ^39^. Overall, all these pieces of evidence support the possibility that the subset of BM-MSCs mostly endowed with immune regulatory function could be the mostly undifferentiated.

In conclusion, we provide evidence that stromal CD40 expression is up-regulated in BM-MSCs after irradiation as a step necessary to restore T cell homeostasis during HSCT. CD40-deficiency in BM stromal cells skews BM T cell activation towards the persistent production of inflammatory molecules that, in turn, impair normal B cell development. These CD40-dependent changes are pathogenetically relevant in human aGVHD. Based on our data, CD40 appears to be a key molecule conferring tolerogenic properties to BM niches.

## Acknowledgments

The authors thank the Conventional and Confocal Microscopy Facility for confocal images acquisition. This work was supported by the Italian Ministry of Health (GR-2013-02355637 to S. Sangaletti); and Associazione Italiana per la Ricerca sul Cancro (Investigator Grant number 22204 to Sabina Sangaletti and 10137 to M.P. Colombo). Barbara Bassani is funded by the FIRC-AIRC (Fondazione Italiana per la Ricerca sul Cancro) fellowship “Guglielmina Lucatello e Gino Mazzega”. The authors also thank E. Grande for administrative support.

## Author contributions

S.S and M.P.C. designed research; B.B. A.P., A.G. P.P. L.B. B.C. M.C. performed the experiments, S.S., I.N. K.J. I.A. provided human biopsies and analysed clinical parameters, C.C. C.T. A.C. analyzed the data. S.S. N.B., C.T. and M.P.C wrote the manuscript.

## Additional information

**Supplemental Information** accompanies this paper.

