## Supplementary material for "CD40 provides immune privilege to the bone marrow hematopoietic niche": Supplemetary Figures

SUPPLEMENTAL METHODS, FIGURES AND TABLES

**METHODS**

**Histology.** In GVHD experiments, histological damage was evaluated in the skin, liver, lung and kidney according to a semiquantitative scoring system based on histopathological analysis of routinely-stained haematoxylin and eosin 4-micra-thick tissue sections. In detail, epidermal necrosis, epidermal inflammatory infiltration, and dermal inflammatory infiltration were analysed in the skin; parenchymal inflammatory infiltration, peri-portal necrosis, and centro-lobular necrosis were assessed in the liver; interstitial inflammatory infiltration, epithelial proliferation, and stromal proliferation were scored in the lung parenchyma; glomerular rarefaction, glomerular necrosis, glomerular inflammatory infiltration, and interstitial inflammatory infiltration were evaluated in the kidney. Each variable was scored according to a four-grade semiquantitative score: 0 (absent), 1 (focal/mild), 2 (multifocal/moderate), 3 (diffuse/intense). The final damage score was calculated for each experimental condition by averaging the scores relative to the same tissues.

**Lentiviral vector construction, virus production and MSC infection.** To construct CXCR4-expressing lentivector, we modified the self-inactivating lentiviral vector pRRLCMVGFPsin-18 [(kind gift of Dr. G. Ferrari {Lotti, 2002 #57}] by replacing the GFP sequence with mouse CXCR4 cDNA (Genescript). A third-generation packaging system (pMDLg/pRRE and pRSV-REV and pMD2-VSV-G and transfer vectors) was used to produce viral particles. Lentiviral stocks were produced in 293T cells by $Ca_3PO_4$ co-trasfection of the four plasmids. Twenty-four hours after transfection, the virus containing supernatant was harvested, filtered and purified by ultracentrifugation as described previously {De Palma, 2002 #60}. The viral titer was calculated by analysis of the expression of CXCR4 by flow cytometry in 293T cells infected with different dilution of virus. A MOI 50 was used to infect MSC.

**Real Time PCR.** Total RNA was extracted by using the Quick RNA micro prep kit (Zymo Research) and subsequently quantified with NanoDrop 2000c Spectrophotometer (Thermo Scientific). MultiScribe-Reverse Transcriptase kit (Applied Biosystems) was used for the reverse transcription assay and the Real-Time PCR was performed using the Taqman Universal PCR Master Mix (Applied Biosystems) according to the manufacturer instructions. Briefly, the master mix, 20 ng of cDNA and the probes of interest were diluted in a total volume of 20 µL and the Real-Time PCR was performed on 7900HT Fast Real-Time PCR System (Applied Biosystems).

Taqman Probes: *Cd40* (*Mm00441891_m1*), *ido-1 (Mm00492590_m1)*, *Nos2* (*Mm00440502_m1*), *pd-l1* (*Mm01208504_m1*), *sparc* (*Mm00486332_m1*).

**IHC and immunofluorescence analyses.** For IHC, Human and Murine BM samples were fixed in 10% buffered formalin, decalcified using an EDTA-based buffer, and paraffin-embedded. Tissue sections (4 μm) were deparaffinized and rehydrated. An antigen unmasking technique was performed using Novocastra Epitope Retrieval Solution (pH 9) in a PT Link Dako pre-treatment module at 98°C for 30 min. Subsequently, the sections were brought to room temperature and washed in PBS. After neutralization of the endogenous peroxidases with 3% H2O2 and Fc-blocking by a specific protein block (Novocastra, UK), the samples were incubated overnight at 4°C with primary antibodies listed in Supplementary Table 2. The immunostaining was revealed by either a polymer detection method (Novolink Polymer Detection Systems Novocastra Leica Biosystems Newcastle Ltd Product No: RE7280-K), and following specific secondary antibodies: horseradish peroxidase (HRP)-conjugated donkey anti-rabbit IgG (H+L)(A16035, Invitrogen) or HRP-conjugated goat anti-rat IgG (H+L) (Vector Lab) secondary antibody and either 3-amino-9-ethylcarbazole (AEC) or 3,3'-diaminobenzidine (DAB) substrate-chromogens.

For co-immunofluorescence, BM and spleen sections were pretreated as detailed above for IHC. The primary antibodies used for this analysis are detailed in Supplementary Table 2. Primary antibody binding was amplified and visualized using Alexa Fluor 568-conjugated goat anti-rabbit IgG (H+L) (A11011, Invitrogen), Alexa Fluor 488-conjugated goat anti-rat IgG (H+L) (A11006, Invitrogen), Alexa Fluor 488-conjugated goat-anti-mouse IgG (H+L) (A11001, Invitrogen), or Alexa Fluor 633-conjugated goat anti- rat IgG (H+L) (A21094, Invitrogen). The slides were counterstained with DAPI Nucleic Acid Stain (Invitrogen Molecular Probes). The samples were then analyzed under an Axioscope A1 optical microscope (Zeiss) and microphotographs were collected using an Axiocam 503 color digital camera (Zeiss). Images were analyzed using Zen2 imaging software.

The quantitative analysis of CD40 and PAX5 in BM biopsies from human samples by immunohistochemistry was performed on whole sections scans using a Leica Aperio CS2 slide scanner and by quantifying the staining by two different software tools. The Nulcear Hub tool (Nuclear v9) of the Image Scope software was used to determine the percentage of PAX5-expressing nuclei over the total cellularity, while the Positive Pixel Count v9 tool was adopted to categorize the CD40-stained slides according to a semi-quantitative score ranging from 0 (absent) to 3 (diffuse/intense).

For double-marker immunofluorescence, the sections were incubated with the following primary antibodies: Rabbit Polyclonal CD40 and Rabbit Polyclonal Granzyme B. The binding of the primary antibodies to their respective antigenic substrates was revealed by Opal Multiplex IHC kit, which allowed for combined immunostainings using antibodies with a same made through tyramide signal amplification. After deparaffinization, antigen retrieval was performed using microwave heating in pH9 buffer and the first primary antibody was incubated. Immunofluorescence labeling was achieved by incubating with a specific secondary antibody followed by the addition of one selected Opal fluorophore and microwave treatment in pH9 buffer. The same procedure was repeated for the second primary antibody using a different Opal fluoprophore and DAPI nuclear counterstain.

SUPPLEMENTAL FIGURES

**Supplemental Figure 1**

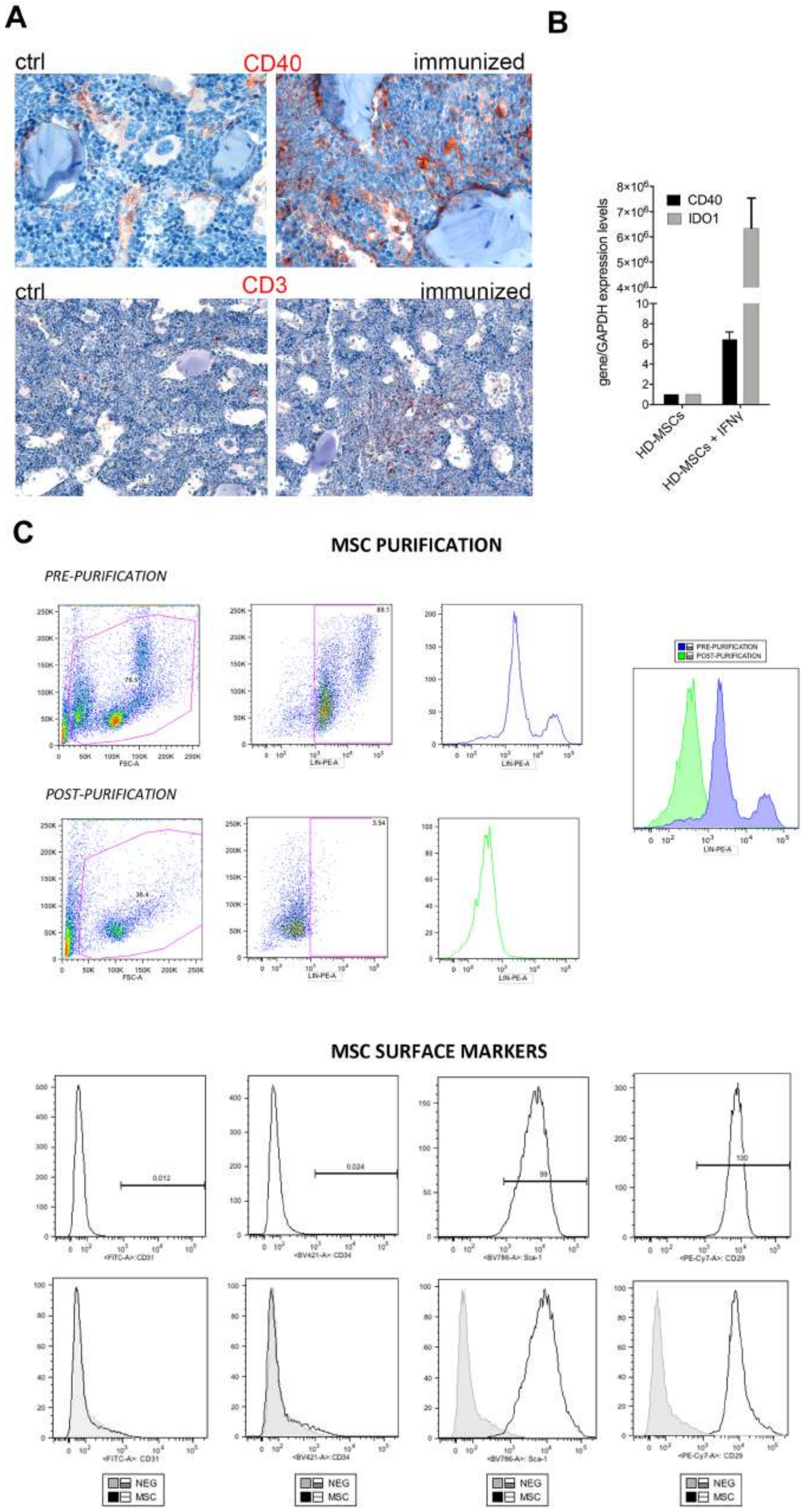


*CD40 expression in BM from autoimmune mice and human BM-MSCs.* **A.** IHC analysis for CD40 and CD3 in BM of mice developing autoimmunity {Tripodo, 2017 #221} and characterized for the expansion of IFNγ+ Teff. **B.** Real-time PCR analysis showing the expression of CD40 and IDO1 in human MSCs obtained from healthy donors (HD-MSCs) upon IFNg stimulation. **C.** Phenotypic characterization of purified mesenchymal stem cells – Following the enzymatic digestion with collagenase 0,2 mg/mL, cell suspension is filtered by using 70 mm cell strainers and incubated with the Lin antibody cocktail that includes CD45, CD11b, CD11c, CD3, GR-1, F4-80, B220, TER119 PE-conjugated. Then cells where labeled with Anti-PE MicroBeads (Miltenyi Biotech) and magnetically separated. Representative dot plots and the relative histograms of the samples pre- and post-purification are shown MSC phenotype was deeper investigated using CD34, CD31, CD29 and Sca-1 antibodies and the expression of the selected surface markers is shown by representative histograms

**Supplemental Figure 2**

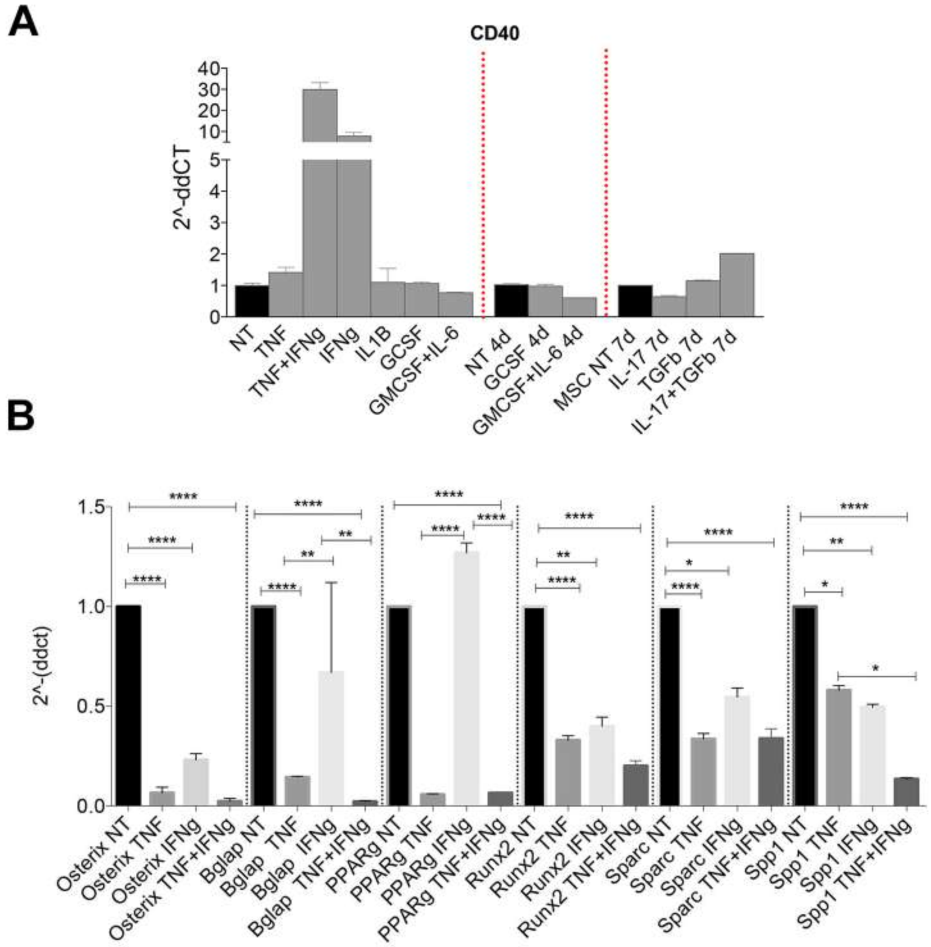


*Expression of BM-MSCs speciation markers upon stimulation with inflammatory cytokines.* **A.** RT-PCR analysis for CD40 expression in BM-MSCs (ex-vivo isolated and in vitro expanded) treated in vitro with IL-1b, G-CSF, GM-CSF+IL-6. **B.** TNF and IFNg stimulation of BM-MSC prevented the differentiation program toward osteo-and adipo- lineages. RT-PCR analyses showed that, after 24hours, the treatment with 10 ng/mL IFNg and 50 ng/mL TNFa decreased the expression of osteoblast and adipocytes differentiation markers on BM-MSC compared with untreated cells. The combination of TNF and IFNg was additive in decreasing the expression of *Spp1* (* $p < 0.05$, ** $p < 0.01$, **** $p < 0.0001$, One-way ANOVA).

**Supplemental Figure 3**

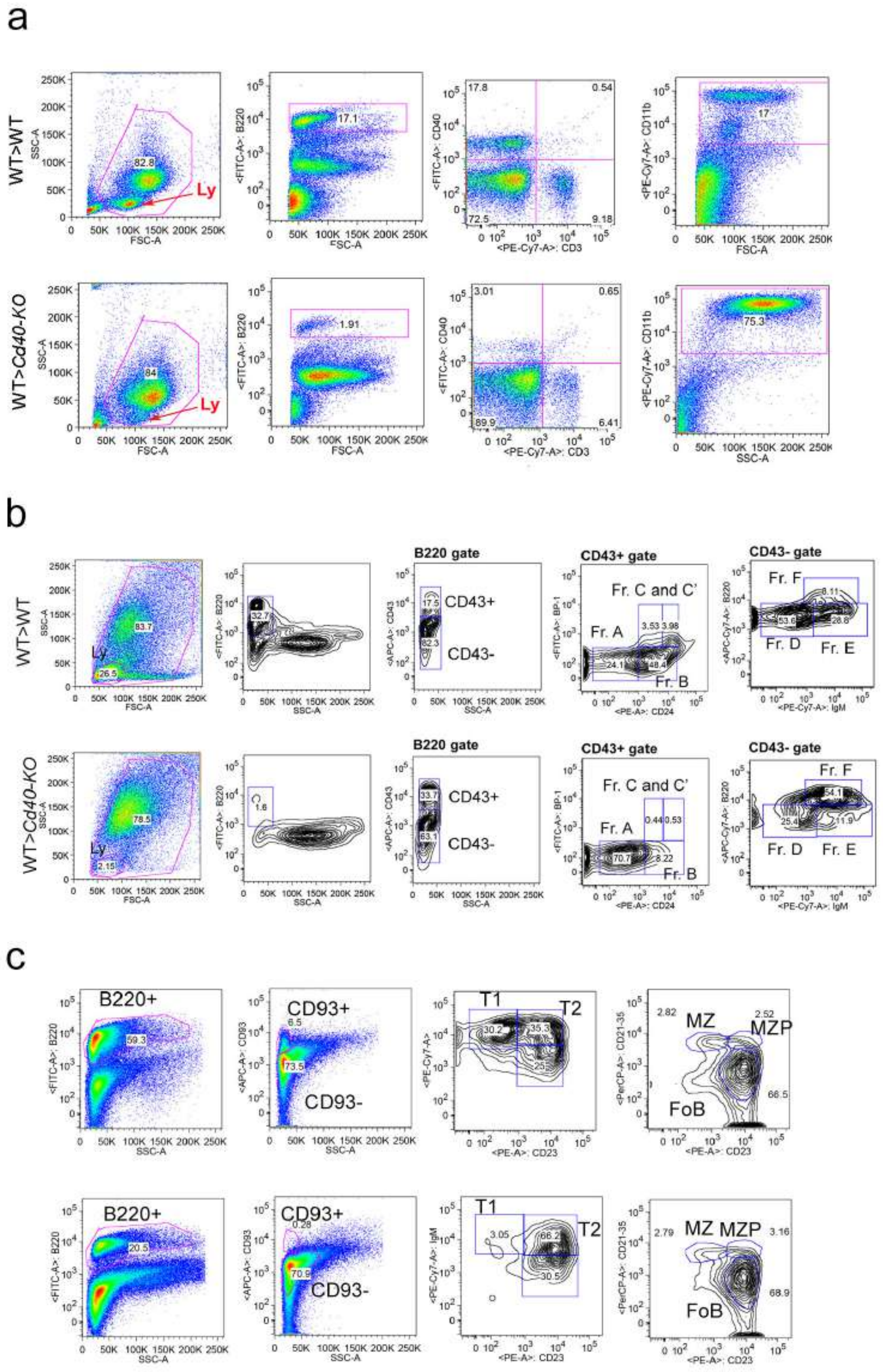


*Gating strategy utilized to characterize B-cell development in mouse BM chimeras.*

**a.** Representative gating strategy for PB FACS analysis showing the frequencies of B220+, CD3 and CD11b+ cells in the PB of WT>*Cd40*-KO BM chimeras compared to WT>WT BM chimeras.
**b.** Representative dot plots showing the gating strategy used to characterize BM B-cell development in chimeric mice. CD43+ B-cell precursors are identified within the B220+ gate. CD43+ cells are further characterized according to the expression of BP-1 and CD24 that allows to

distinguish among the different pre-pro and pro-B fractions (A, B, C and C'). Within the CD43- population the expression of IgM and B220 identified the fraction D, E and F **c** representative dot plots showing the different B-cell populations maturating in the spleens of WT>WT and WT> *CD40-KO* BM chimeras. Splenic immature B220+CD93+ B-cell were divided into transitional T1, T2, and T3 cells based on their expression of CD23 and IgM (T1 = IgM + CD23-, T2 = IgM + CD23+, T3 = $IgM^{low}$CD23+).

**Supplemental Figure 4**

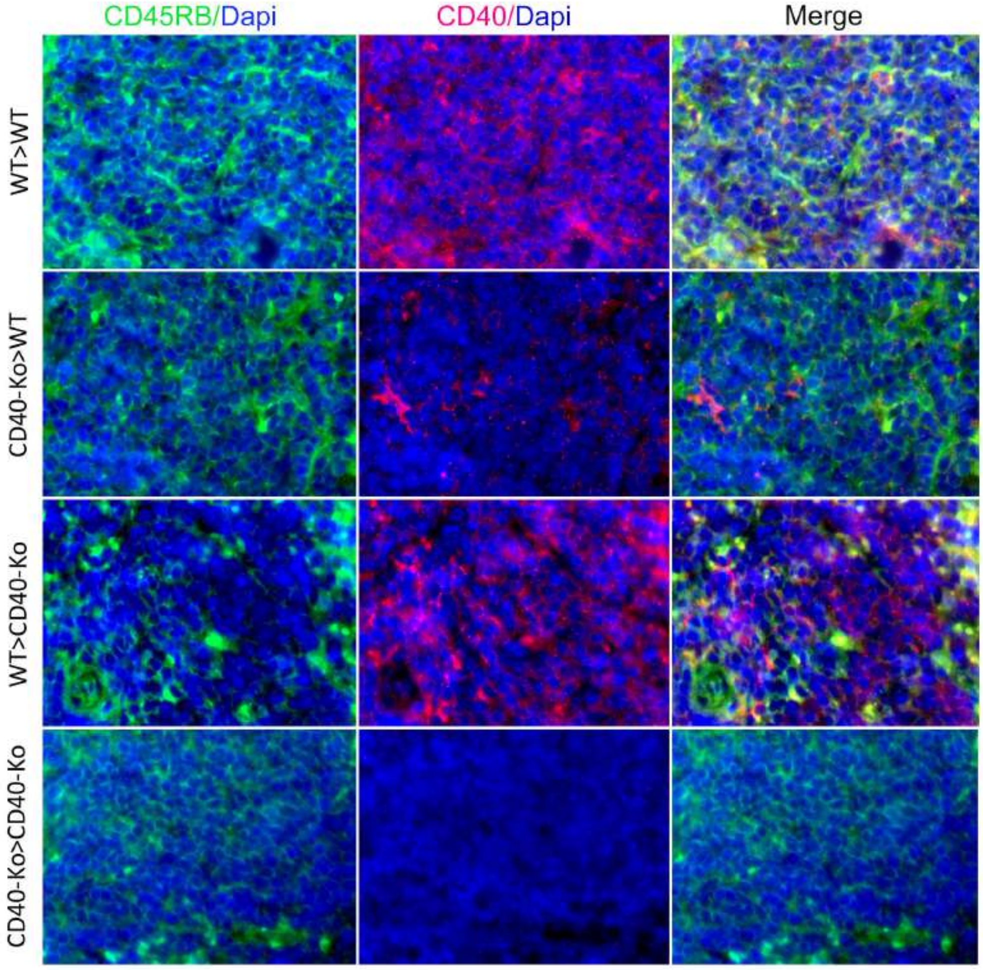


*Detection of B220+CD40+ B-cells in Cd40-KO recipient mice* Representative IF microphotographs of spleen sections from different chimeric mice showing the presence of B220+(green signal)CD40+(red signal) cells in the spleen of *WT>WT* and *WT>Cd40-KO* chimeras. In *Cd40-KO>WT* chimeras CD40 expression is confined to scattared CD45RB B220-negative stromal elements while *Cd40-KO>Cd40-KO* controls have no detectable CD40 expression (Original magnification x400).

**Supplemental Figure 5**

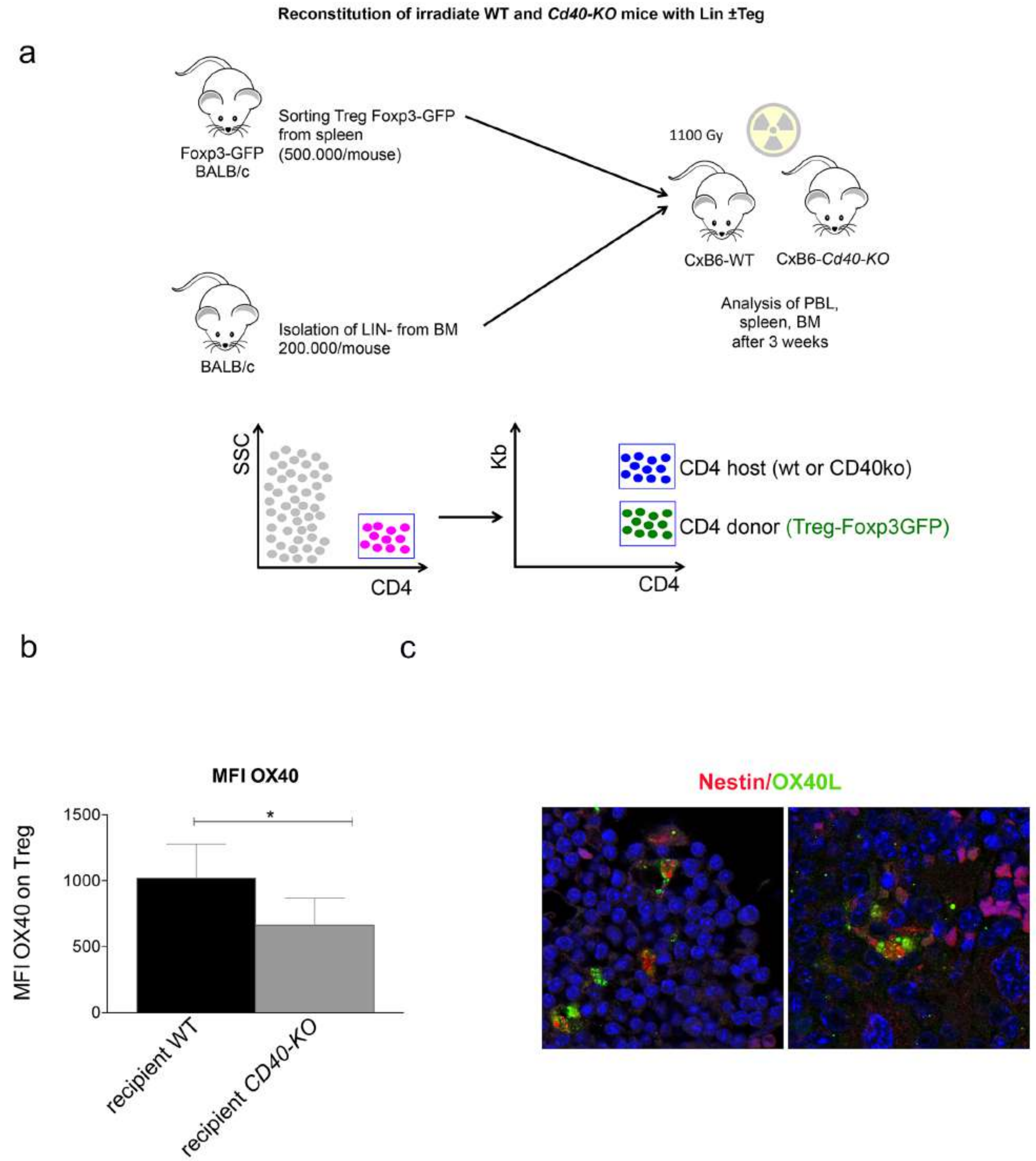


*Reduced OX40 MFI on Treg and increased OX40L expression in the BM microenvironment of CD40-KO recipents.* **a.** Schematic representation of BMT experiments in which Lin- cell were co-injected with Treg cells into lethally irradiated WT and *Cd40-KO* mice. The use of CxB6 F1 mice (either WT or *Cd40-KO*) as recipients allowed discriminating donor and host Treg, being donor Treg H-2Kb negative. b. Cumulative data display the significantly reduced MFI of OX40 in *Cd40-KO* compared to WT recipients (*p<0.05; n=10/group; Student *t* test). IF showed the co-localization between nestin (BM-MSC marker, red) and OX40L (green).

**Supplemental Figure 6**

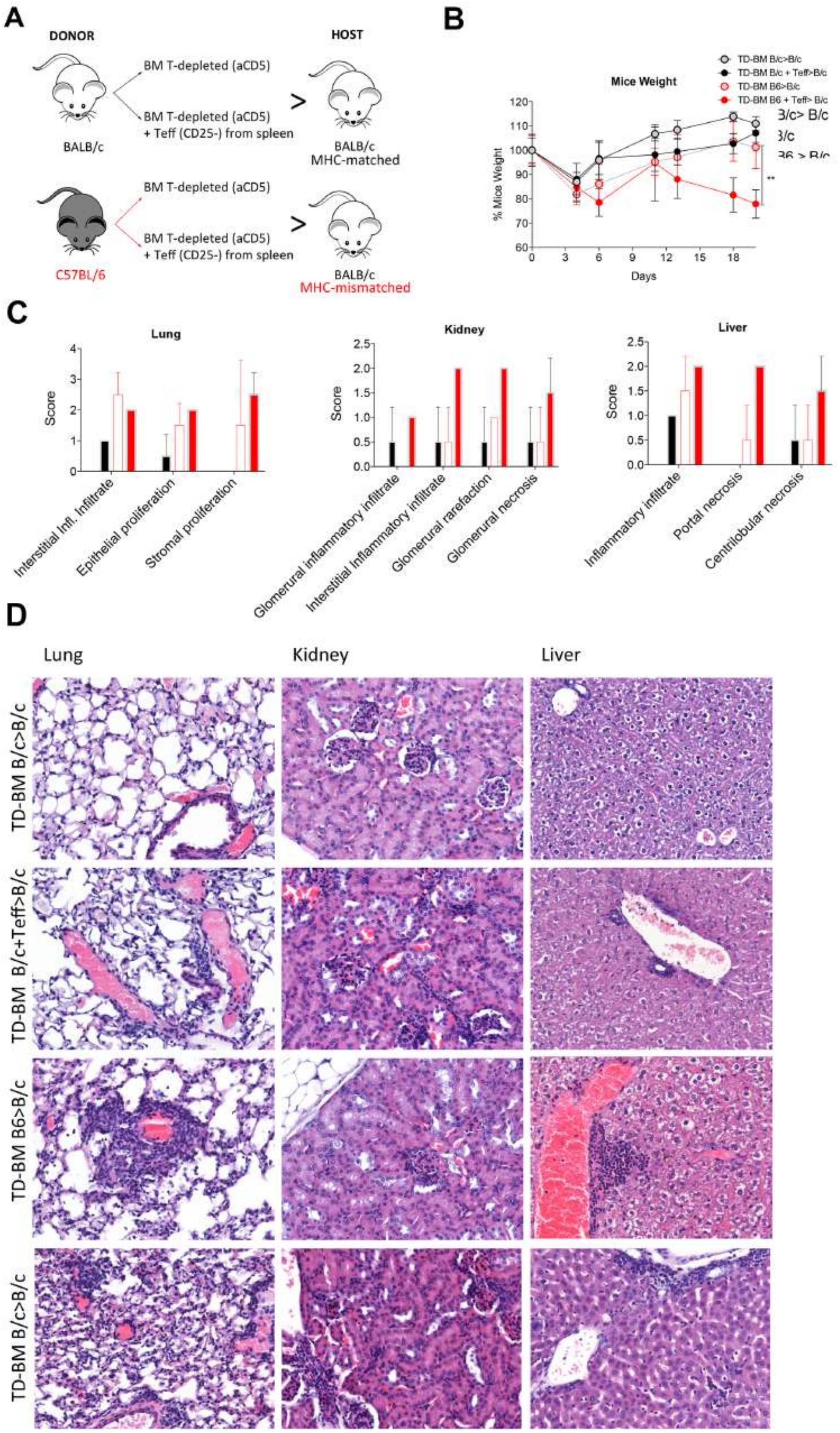


*Histological features of aGVHD mouuse models.* a. Schematic representation of the BMT experiment performed to obtain MHC-mismatched (modeling aGVHD) or non-MHC-mismatched BM chimeras. **b** Weight loss in aGVHD at day 21 after BMT in the specified allogeneictransplanted and control animals (n= 12 per group). **p < 0.005, compared using Student's *t* test. **c**. Histological score for lung, liver, kidnay and skin damage in aGVHD mice (TD-BM+Teff B6>B/c) and control groups. **d.** Representative microphotographs detailing the different extent of immune infiltration and associated histological damage in the lung, kidney and liver parenchyma of mice belonging to the four groups. No significant immune cell infiltration is detected in the tissues of TD-BM B/c>B/c mice; slight inflammatory infiltration mainly confined to peri-vascular areas and without signs of parenchymal damage characterizes TD-BM B/c+Teff>B/c mice; Focally severe inflammatory infiltration associated with evidence of parenchymal damage is detected in TD-BM B6c>B/c mice samples; Multifocal or diffuse immune infiltration determines overt disarrangement of the histological architecture in the three organs of TD-BM B6+Teff>B/c mice.

**Supplemental Figure 7**

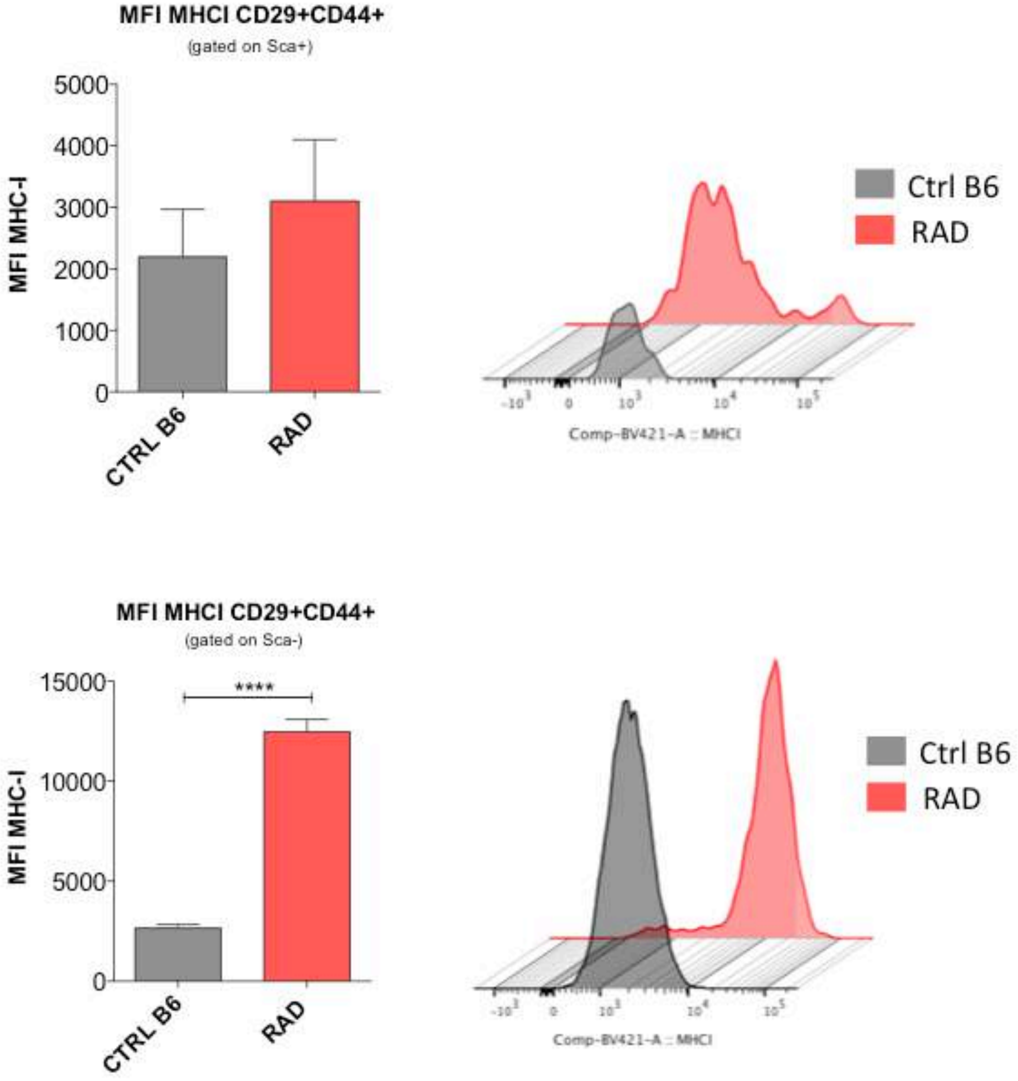


*MHC-I expression on Sca+ and Sca- BM-MSCs (CD29+CD44+) upon lethal irradiation.*
Cumulative data displaying the increase MFI of MHC-I in irradiated compared to non irradiated (control) mice. Representative histograms are also shown. ****$p < 0.0001$, compared by Student's *t* test.

**Supplemental Figure 8**

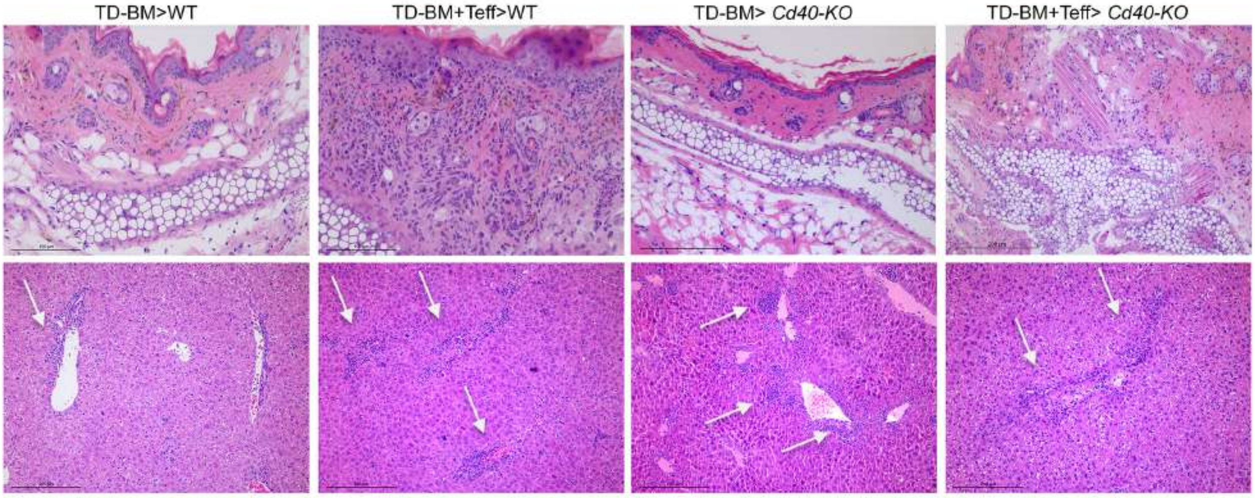


*Histological features of aGVHD in Cd40-KO recipients.* Representative microphotographs detailing the different extent of immune infiltration and associated histological damage in the skin and liver parenchyma of mice. Focally inflammatory infiltration associated with evidence of parenchymal damage is detected in TD-BM B6>B/c mice samples; Multifocal or diffuse immune infiltration characterized the liver of all Cd40-KO recipents, also in the absence of Teff (TD-BM> *Cd40-KO*).

**Supplemental Figure 9**

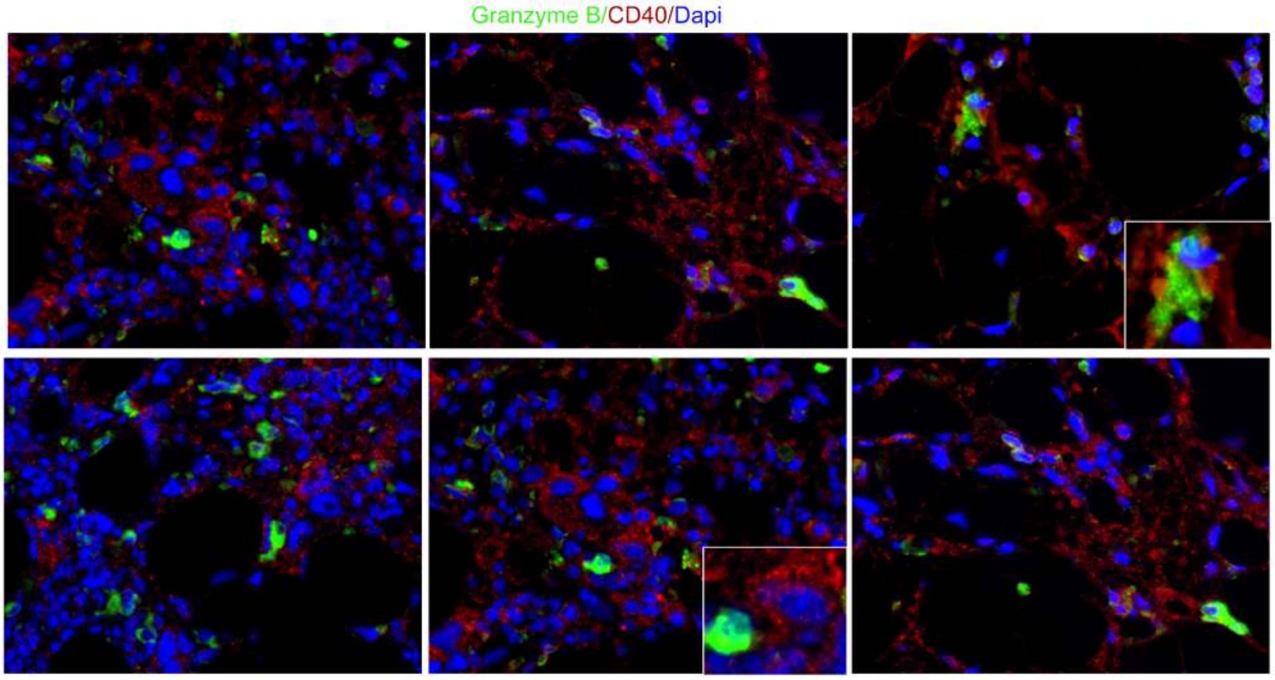


*CD40+ stromal cells are co-localized with Granzyme B granules in aGVHD BOM.* IF analysis showed the co-localization between Granzyme B (green) and residual CD40+ BM-MSCs (red) in aGVHD patients  previously characterized fir residual PAX5 staining (see Figure 7) . The insets are showing the close contact between granzyme + (green) and CD40+ (red) elements.
