## Supplementary Tables for "CD40 provides immune privilege to the bone marrow hematopoietic niche"

**Supplemental Table 1. Patients characteristics.**

| Patient | day of biopsy post-transplant | underlying disease | age at transplantation/sex | ATG | immunosuppression | aGVHD | cGVHD | IHC Pax5 | IHC CD40 |
|---|---|---|---|---|---|---|---|---|---|
| #1 | d28 | AML | 59/m | x | Cyclosporine, Cellcept | Grade III | | <1% | 1 (myelo) |
| #2 | d26 | AML | 45/m | | Cyclosporine, MTX | Grade III | | <1% | 0 |
| #3 | d25 | AML | 57/f | x | Cyclosporine, Cellcept | | | 6% | 1 (myelo) |
| #4 | d21-28 | T-NHL+tAML | 71/f | x | Cyclosporine, Cellcept | | | 21% | 2 (myelo + stroma) |
| #5 | d21 | AML | 72/m | x | Cyclosporine, Cellcept | | Grad III | <1% | 0 |
| #6 | d24 | Ph+ ALL | 58/f | | Cyclosporine, MTX | Grade I | | 5% | 2 (myelo) |
| #7 | d28 | AML | 43/f | | Cyclosporine, MTX | Grade II | | 3% | 1 (myelo) |
| #8 | d28 | AML | 62/f | x | Cyclosporine, Cellcept | | | 15% | 2 (myelo + stroma) |
| #9 | d28 | AML | 57/m | x | Cyclosporine, Cellcept | Grade II | | 8% | 1 (myelo + stroma) |
| #10 | d22 | AML | 58/f | x | Cyclosporine, Cellcept | | | 12% | 1 (myelo + stroma) |
| #11 | d28 | ALL | 23/m | | Cyclosporine, MTX | Grade II | | <1% | 1 (myelo) |
| #12 | d28 | AML | 60/m | x | Cyclosporine, Cellcept | Grade II | | <1% | 1 (myelo) |

**Supplemental Table 2. List of antibodies.**

| Fluorescence | Antigen | Clone | Manufacturer |
|---|---|---|---|
| **BM ANALYSIS** | | | |
| **APC-eFluor 780** | **B220** | RA3-6B2 | eBioscience |
| **APC** | **CD43** | S7 | BD |
| **PECy7** | **IgM** | R6-60.2 | BD |
| **PE** | **CD24** | 30-F1 | BD |
| **PE** | **B220** | RA3-6B2 | BD |
| **FITC** | **BPI** | F6354 | eBioscience |
| **SPLENIC B-CELL COMPOSITION** | | | |
| **FITC** | **CD43** | eBioR2/60 | eBioscience |
| **APC-Cy7** | **B220** | RA3-6B2 | Biolegend |
| **APC** | **CD93** | AA4.1 | eBioscience |
| **PE** | **CD23** | B3B4 | eBioscience |
| **FITC** | **IgD** | 11-26c | eBioscience |
| **PerCP** | **CD21/35** | 7E9 | Biolegend |
| **APC** | **CD3** | 145-2C11 | BD |
| **PE** | **CD11c** | N418 | Tonbo Biosciences |
| **Pe-Cy7** | **CD11b** | M1/70 | Tonbo Biosciences |
| **APC -e-fluor780** | **Gr-1** | RB6-8C5 | eBioscience |
| **T-CELL ANALYSIS** | | | |
| **FITC** | **CD25** | eBio7D4 | eBioscience |
| **FITC** | **TNFα** | MP6-XT22 | BD |
| **PE** | **OX40** | OX86 | Biolegend |
| **PerCPCy5.5** | **Foxp3** | FJK-16S | eBioscience |
| **PE-Cy7** | **CD4** | RM4-5 | Tonbo Biosciences |
| **APC** | **IFNγ** | XMG1.2 | eBioscience |
| **APC-eFluor780** | **CD45.2** | L104 | Biolegend |
| **APC-eFluor780** | **CD45.1** | A20 | eBioscience |

| Fluorescence | Antigen | Clone | Manufacturer |
|---|---|---|---|
| **BM-MSC ANALYSIS** | | | |
| **FITC** | **CD45** | 30-F11 | Tonbo Biosciences |
| **FITC** | **CD31** | 390 | eBioscience |
| **FITC** | **TER119** | Ter119 | Invitrogen |
| **FITC** | **CD34** | RAM34 | BD |
| **FITC** | **cKIT** | 2B8 | eBioscience |
| **APC** | **SCA-1** | D7 | BD |
| **APC-Cy7** | **CD44** | IM7 | BD |
| **BV421** | **MHC-I** | AF6-88.5 | BD |
| **BV605** | **CD73** | TY/11.8 | Biolegend |
| **BV711** | **MHC-II** | M5/114.15.2 | BD |
| **PE** | **CD40** | 3/23 | BD |
| **BV421** | **CD40** | 3/23 | BD |
| **PE-Cy7** | **CD29** | eBioHMb1-1 | eBioscience |

| Antigen | Type | dilution | Manufacturer |
|---|---|---|---|
| **Immunohistochemistry (IHC)** | | | |
| **PAX-5 (ab109443)** | monoclonal anti-human | 1:1000 | Abcam |
| **CD40 (ab13545)** | polyclonal anti-human and mouse | 1:100 | Abcam |
| **iNOS (ab15323)** | polyclonal anti-mouse | 1:100 | Abcam |
| **PD-L1 (D5V3B)** | monoclonal anti-mouse | 1:200 | Cell Signaling |
| **OX40L (RM134L)** | monoclonal anti-mouse | 1:100 | BD |

| Co-immunofluorescence | | | |
|---|---|---|---|
| **B220 (9F14)** | monoclonal anti-mouse | 1:50 | Genetex |
| **CD40 (ab13545)** | polyclonal anti-human and mouse | 1:100 | Abcam |
| **OX40L (RM134L)** | monoclonal anti-mouse | 1:100 | BD |
| **CD146 (ab49492)** | monoclonal anti-human | 1:25 | Abcam |
| **anti-nestin PE-conjugated** | polyclonal | 1:100 | Invitrogen |
